# Substrate tunnel regulates substrate specificity switching in hyaluronan synthase

**DOI:** 10.1101/2024.10.24.620076

**Authors:** Kaiyi Zhu, Yilei Han, Yupei Jian, Guoqiang Jiang, Diannan Lu, Jianzhong Wu, Zheng Liu

## Abstract

Hyaluronic acid (HA), a ubiquitous linear polysaccharide in the extracellular matrix of vertebrates and the capsule of certain pathogenic bacteria, is characterized by its unique structure of alternating β-1,3-*N*-acetylglucosamine and β-1,4-glucuronic acid units. HA is synthesized by Class I hyaluronan synthase (HAS) in the majority of organisms, but the molecular mechanisms determining the alternating sequence of HA remain unclear. Here, we demonstrated that HA synthesized *in vitro* using *Streptococcus equisimilis* HAS maintained its alternating sequence, regardless of substrate ratios and concentrations. While the active pocket lacks intrinsic selectivity, a distal allosteric regulation from the transmembrane tunnel is revealed. The lingering HA terminals in the active pocket ensure lower binding energies for alternating substrates through Coulombic repulsion and competitive binding. Moreover, the dynamic C-terminal loop at the substrate tunnel entrance reduces transport distance for alternating substrates. Our findings indicate that HA-substrate interactions and enzyme dynamics in the substrate tunnel collectively govern the alternating substrate specificity of HAS. This study provides the first molecular insight into precise substrate specificity control within a single active pocket during template-independent polymer synthesis.

## Introduction

Hyaluronic acid (HA) is a linear glycosaminoglycan featured by alternating β-1,3-*N*-acetylglucosamine (GlcNAc) and β-1,4-glucuronic acid (GlcUA) unit. As a critical component of vertebrate extracellular matrix (ECM), HA is involved in numerous physiological and pathological processes such as cell differentiation, tissue repair, inflammation, tumorigenesis, cancer progression, metastasis and etc^1, 2^. In certain bacterial pathogens, such as Group A *Streptococcus* and *Pasteurella multocida*, HA forms a capsule that acts as a camouflage to facilitate immune evasion and enhance virulence^3^. These processes are triggered by the recognition of HA’s unique alternating sequence by cell surface receptors including CD44, TLR4, and RHAMM^4, 5, 6^

Class I hyaluronan synthase (HAS) catalyzes HA synthesis in the majority of organisms. In this process, HA is elongated at the periplasmic glycosyltransferase (GT) domain and transported through the HAS transmembrane (TM) tunnel, thereby enabling processive synthesis^7^. Remarkably, Class I HAS exhibits a single active pocket, but regulates the alternating glycosyl transfer of two substrates, UDP-GlcNAc (UGN) and UDP-GlcUA (UGA)^7^. It provides the first and only example that challenges the “one pocket, one sugar” principle^7^. While the structures of HAS-substrate complexes have been predicted using molecular dynamics simulation^13^ and resolved by cryo-EM recently^8^, the mechanism underpinning the alternating selection of two substrates remains unknown.

This study is devoted to a molecular-level understanding of the alternating substrate specificity of single-pocket *Streptococcus equisimilis* HAS (SeHAS), using molecular simulations and mutagenesis experiments. SeHAS demonstrated its alternating substrate selectivity *in vitro*, regardless of the dimerization of SeHAS. The effects of conformational dynamics of the active site, substrate tunnel, and the distal transmembrane tunnel on substrate selectivity are examined, respectively. The collaboration of HA-substrate interactions and HAS conformation dynamics on the alternating substrate selectivity is displayed.

## Results

### SeHAS maintained alternating substrate specificity *in vitro*

Liposomes embedded with SeHAS were prepared as *in vitro* catalysts by ultrasonication following cell disruption. Transmission electron microscopy (TEM) and dynamic light scattering (DLS) measurements confirmed the liposome morphology, and sodium dodecyl-sulfate polyacrylamide gel electrophoresis (SDS-PAGE) demonstrated the loading of full-length HAS in the liposomes. HA was synthesized within these liposomes using UDP-GlcUA and UDP-GlcNAc as the HAS substrates. The polymer was subsequently transported into a vesicular lumen^9^ (Extended Data Fig. 1A-D).

As the polymerization reaction progressed, gel filtration chromatography (GFC) detected the elongation and accumulation of HA within two hours, which could be specifically hydrolyzed by hyaluronidase (HAase). HA synthesized *in vitro* exhibited a molecular weight (*M_w_*) approximately twice that of the cellular product, around 2 MDa, and demonstrated a more uniform molecular weight distribution, with a polydispersity index (*M_w_/M_n_*) close to 1. Moreover, the molecular weight of HA remained consistent across different substrate concentrations and ratios (Extended Data Fig. 1E-I).

At a 1:1 molar ratio of substrate feeding, anion-exchange high-performance liquid chromatography (AEX HPLC) confirmed that substrate consumption was balanced at 1:1. Furthermore, substrate hydrolysis was undetectable, indicating that the substrates were utilized in a 1:1 ratio for the polymer synthesis (Fig. 1A, Supplementary Fig. 1). The 600 MHz proton nuclear magnetic resonance (^1^H NMR) spectrum revealed the ratio of hydrogen atoms on the anomeric carbons, C2 of GlcUA, and the methyl group of GlcNAc to be 2:1:3, indicating that the two monosaccharides were incorporated into HA at a 1:1 ratio (Fig. 1B). Chondroitinase ABC (ChABC) specifically catalyzes the endolytic cleavage of HA, resulting in the production of the unsaturated disaccharide ((4-deoxy-α-L-*threo*-hex-4-enepyranosyluronic acid)-β-1,3-(*N-*acetylglucosamine), ΔHA2) and terminal monosaccharide (4-deoxy-α-L-*threo*-hex-4-enepyranosyluronic acid, ΔUA) as end products^10^ (Supplementary Fig. 2A). GFC confirmed the complete digestion of HA by ChABC (Supplementary Fig. 2B-C), and subsequent electrospray ionization mass spectrometry (ESI-MS) analysis identified ions matching ΔHA2 and ΔUA as the end products. ESI-MS/MS further substantiated the alternating structure of these disaccharides (Fig.1C-D, Supplementary Fig. 3), validating that HA synthesized *in vitro* accurately replicated its natural alternating sequence.

**Fig. 1.**
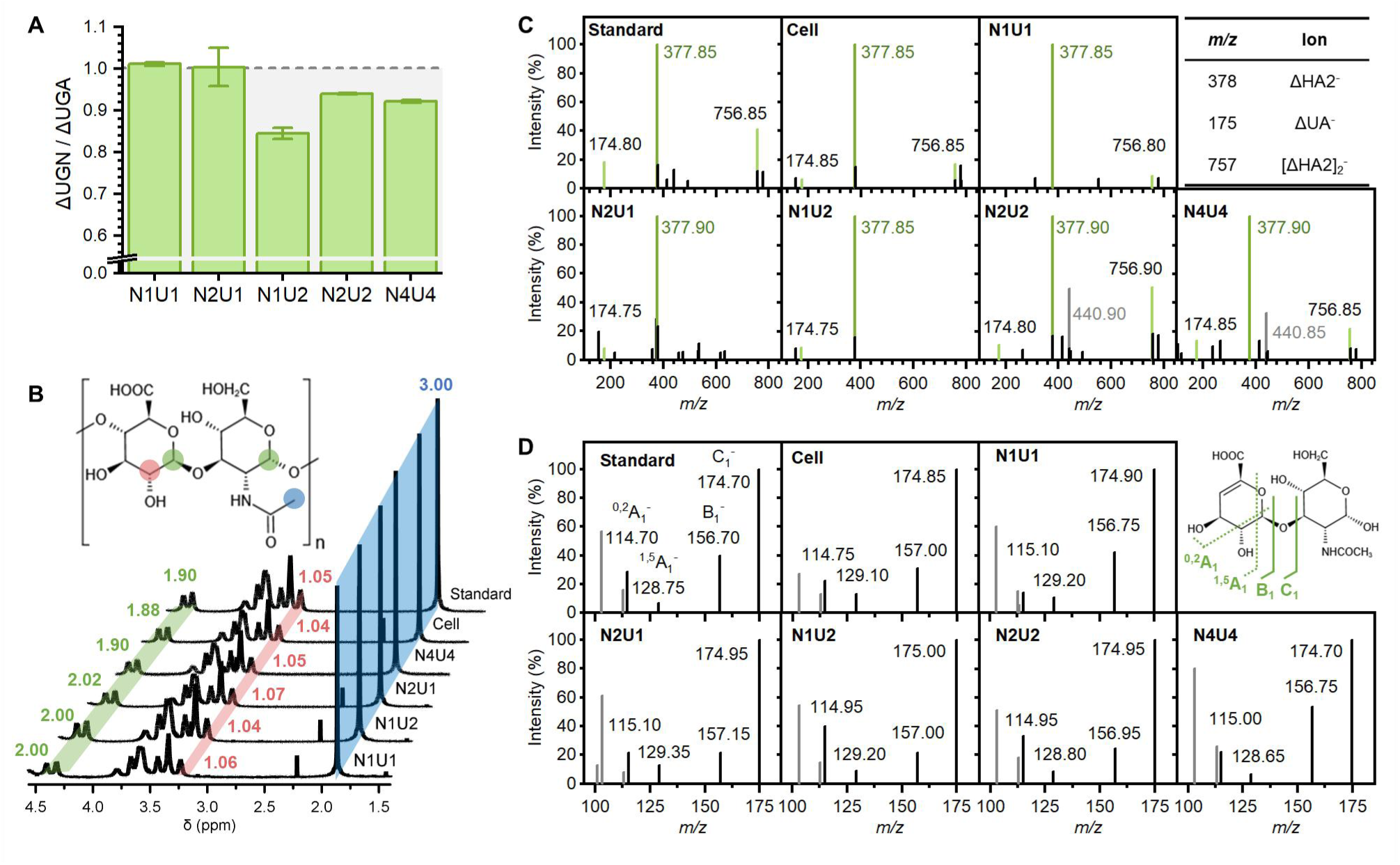
The alternating sequence is preserved for HA synthesized *in vitro*. (A) Molar ratios of substrate consumption. (B) ^1^H NMR spectrum of the HA product. Peak integrals are labeled and normalized with GlcNAc methyl set to 3.00. (C) ESI-MS analysis of HA degraded by chondroitinase ABC. The *m/z*=441 ions are complexes of ΔHA2 and ΔUA. (D) ESI-MS/MS analysis of the *m/z*=378 ion at a collision energy of 20 V. Unlabeled gray peaks are from GlcNAc units. “N”: UGN, “U”: UGA, numbers indicate original concentrations (mmol/L).

Notably, the alternating structural pattern of HA synthesized *in vitro* was unaffected by substrate concentrations and stoichiometric ratios. Despite deviations from the 1:1 substrate consumption ratio under high substrate concentrations or an excess of UGA, ^1^H NMR and ESI-MS confirmed the consistence of 1:1 monomeric unit composition and alternating sequence of high-*M_w_* HA product, respectively (Fig.1, Supplementary Fig.3). Hence, we conclude that the alternating structure of HA must be controlled by the substrate specificity of SeHAS.

### Alternating substrate specificity of SeHAS was governed by a single active pocket

The active unit of SeHAS has been disputed between monomers^11^ and dimers^12^, with the latter suggesting that the alternating linkage of two substrates is achieved by two active pockets. Thus, this study began with the determination of the minimal functional unit for specificity regulation. Both AlphaFold2-Multimer^13^ prediction and HADDOCK^14^ protein-protein docking suggested a “back-to-back” orientation of the two active pockets in HAS dimers (Fig. 2A). The HADDOCK-predicted dimer, reasonably oriented in the membrane as indicated by PPM^15^, was used for further molecular dynamics (MD) simulations, with its TM domains inserted perpendicularly into the bilayer and both GT pockets facing the solution on the same side. Long-term coarse-grained simulations (4∼6 μs) revealed that the HAS dimers maintained a “back-to-back” orientation (Fig. 2B). Notably, negatively charged phospholipids played an important role in maintaining the structure and orientation of HAS dimers. In the PE/PG membrane without divalent cardiolipin (CDL2), GT pockets were partially embedded in the membrane, while in the PE membrane without negatively charged phospholipids, the dimer was completely denatured. Although HAS dimers correctly folded and inserted into both CL and IM membranes, the TM helices in CL membrane exhibited tighter binding and a more “upright” insertion, thereby stabilizing the catalytic pockets in solution. Moreover, all-atom MD simulations identified residues K31, K34, Y37, F39, L41 at the dimer interface (Fig. 2C), aligning with reported cross-linking experiments^12^. Given the “back-to-back” orientation of the active pockets, we conclude that HAS would not acquire expanded active pockets or additional active sites, even in its dimer form. Therefore, the alternating specificity of HAS must be solely achieved through a single active pocket.

**Fig. 2.**
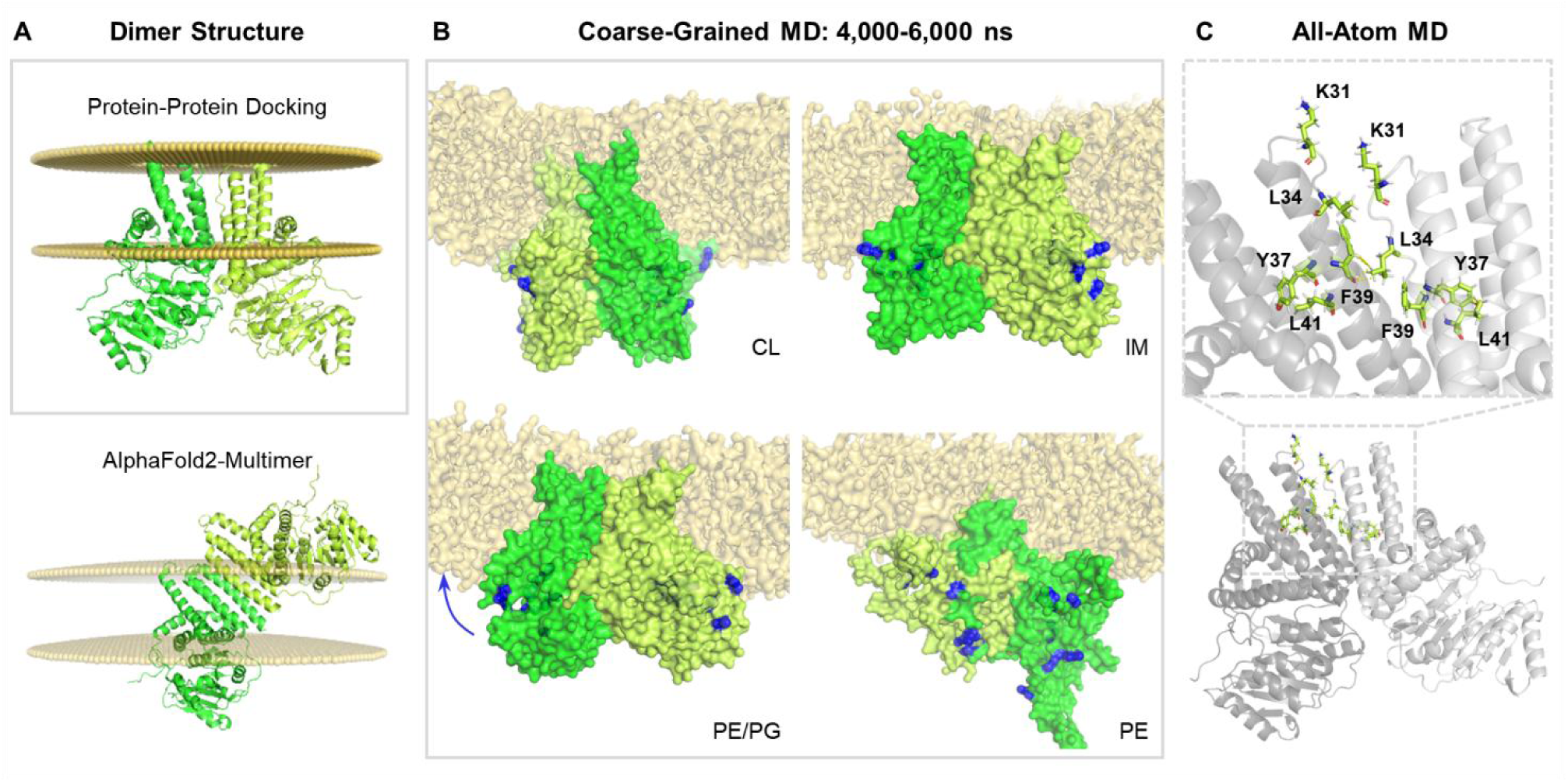
The “back-to-back” orientation of a HAS dimer. (A) Dimer structure predicted by protein-protein docking and AlphaFold2-Multimer. Yellow spheres represent the lipid membrane. (B) Dimer conformations in different membranes. Blue residues represent the GT pocket (E76, D159, D161, D259, D260, R298, W299, T412, K415). (C) The HAS dimer interface predicted by all-atom MD simulation.

### The active pocket failed to discriminate the alternating substrates

In order to discern potential factors influencing substrate selectivity, we compared the binding and transport energies of different substrates within the static active pocket. The SeHAS structure predicted by AlphaFold2^16^ revealed a smooth connection between the TM tunnel and the substrate tunnel in the GT domain, enabling the transport of substrates and HA disaccharides, thus representing a characteristic conformation during HA synthesis (Extended Data Fig. 2).

We analyzed substrate binding within the active pocket using molecular docking, starting with HA docking in the TM tunnel (Fig. 3A). We used two distinct HA tetrasaccharides with terminal GlcUA or GlcNAc, designated as U-HA4 and N-HA4, respectively. These carbohydrates occupy fixed binding sites along the TM tunnels, consistent with previously reported cryo-EM structures^17^. So the positions of two HA terminals varied by one saccharide unit, with GlcUA and GlcNAc located at the entrance of TM tunnel and just outside the entrance, respectively. Subsequent substrate docking revealed that HA terminals directed substrate positioning through hydrogen bonding interactions (Fig. 3B). However, the two substrates adopted similar binding conformations and overlapped in their binding sites, with only slightly lower binding energy (<1 kcal/mol) for alternating substrates.

**Fig. 3.**
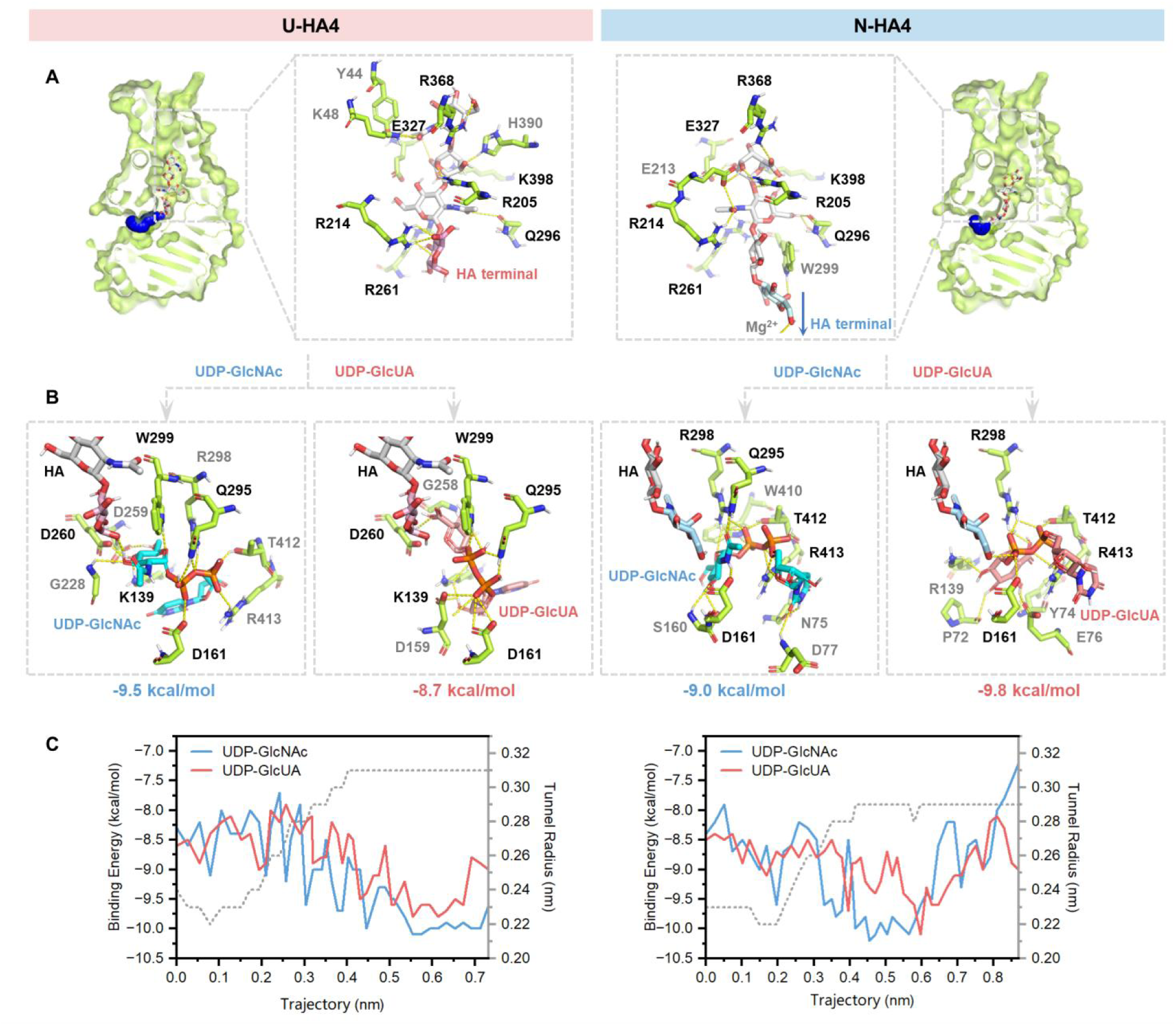
Indistinguishable substrate affinity in the static active pocket of SeHAS. (A) HA binding sites and substrate tunnels. The blue spheres represent the substrate tunnels, with the positions of HA terminal C1 marking as the end point of tunnels. (B) Binding sites and binding energies of different substrates. (C) Transport trajectories and binding energies of different substrates, with the centroids of substrates coinciding the end points of substrate tunnels.

Tunnel analysis (based on Caver Web 1.0^18^) indicated that there was only a single substrate tunnel in SeHAS (Fig. 3A). It transported both substrates with a low binding energy, and the energy variation along the transport path was insignificant (<1 kcal/mol) (Fig. 3C). These indicate that the HAS active pocket does not directly confer alternating selectivity.

### Allosteric regulation of active pocket from TM tunnel

In addition to enzyme-substrate interactions, enzymatic products have the potential to act as allosteric effectors for feedback regulation^19, 20^. To examine the possible allosteric regulation of the TM tunnel, we employed the network-based “Ohm” method^21^ and analyzed the allosteric correlation between the GT pocket and seven glycosyl binding sites along the TM tunnel (Extended Data Fig. 3). Our analysis revealed notable allosteric coupling between the TM outlet and the GT pocket, facilitated through the N-terminal helix. Moreover, conserved catalysis-relevant residues, such as the GDD motif for catalysis and D159 in the DxD motif for substrate or metal ion binding^22^, emerged as significant allosteric hotspots (Fig. 4A-C).

**Fig. 4.**
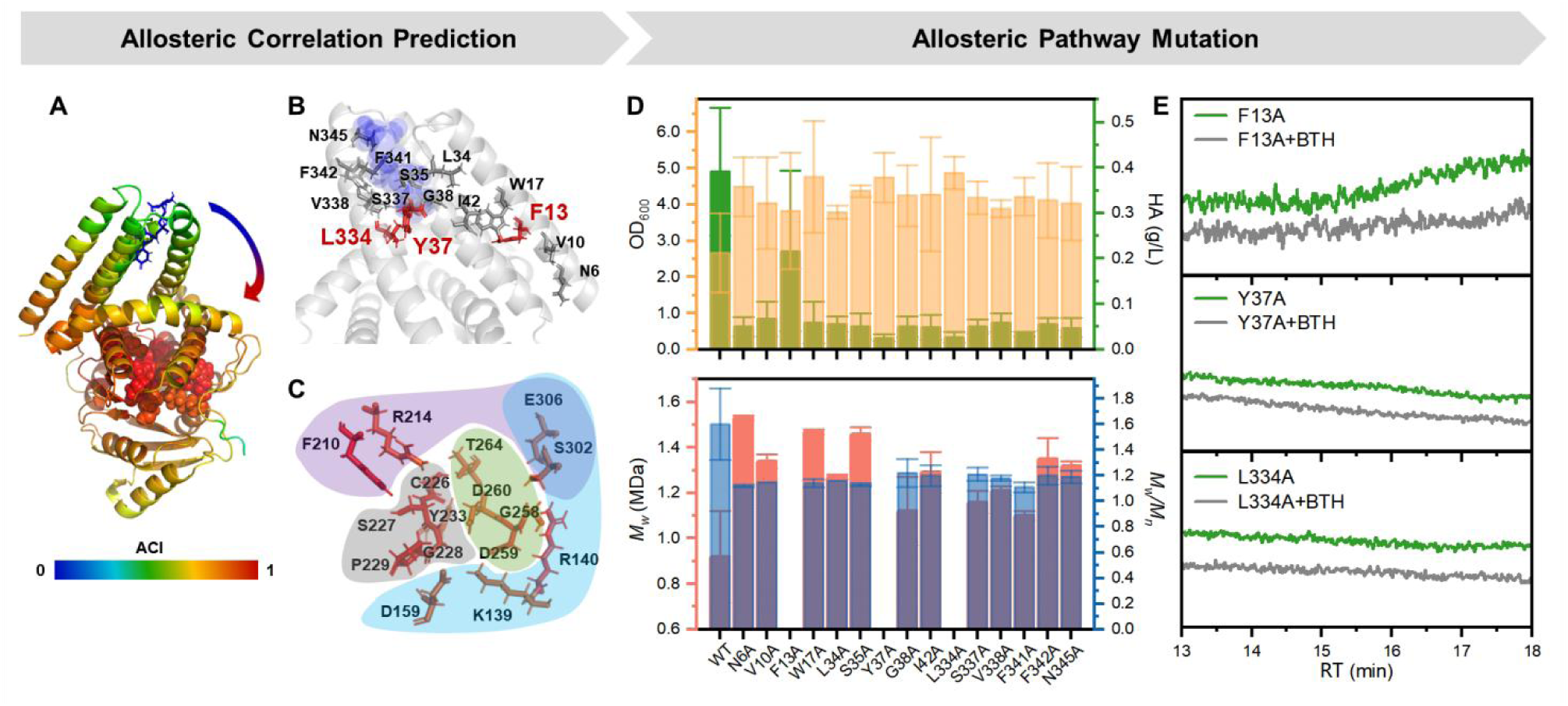
Allosteric correlation between TM outlet and GT pocket. (A) Allosteric coupling intensity (ACI) of HAS residues, sugar at TM outlet as allosteric effector (blue sticks). The red spheres represent allosteric hotspots. (B) Allosteric pathway. (C) Conserved catalysis-relevant residues as allosteric hotspots. The purple region highlights HA binding, the blue region represents substrates binding, the green region is responsible for the catalysis, and the gray region determines the pocket scaffold. (D) The effects of alanine scanning mutagenesis along the allosteric pathway on HA yield, OD_600_, HA *M_w_* and *M_w_/M_n_*. (E) No macromolecular products were detected by GFC analysis for F13A, Y37A, and L334A mutations.

Alanine scanning mutagenesis along the allosteric pathway resulted in decreased HA yield and a metabolic flux shift towards cell growth, leading to higher OD_600_ values (Fig. 4D). Nearly all mutations exhibited an increase in *M_w_*, with the highest observed nearly doubling that of the wild type (WT), reaching *M_w_*=1.5 MDa for N6A (Fig. 4D). Conversely, F13A, Y37A, and L334A failed to synthesize high-molecular-weight HA (Fig. 4E). Among these, F13A retained approximately 50% of the total HA yield (Fig. 4D), suggesting its role in modulating *M_w_* independently of enzymatic activity. Notably, F13 was located at the periphery of the TM tunnel and did not directly interact with HA (Fig. 4B). While previous studies have noted the impact of extensive truncation of the TM domain^23^ and complete replacement of the N-terminal helix^24^ on HA synthesis, our findings represent the first demonstration of the refined regulation of HA yield and molecular weight by an individual peripheral residue in the TM tunnel (F13). In conclusion, our allosteric analysis and mutagenesis experiments reveal that the TM tunnel outlet exerts a distal allosteric regulation on the active pocket, underscoring the necessity of exploring protein dynamics for specificity control.

### HA terminals modulated lower binding energy for the alternating substrate

We employed all-atom MD simulations to explore the influence of HA binding on the conformation and substrate specificity of active pockets. Four HA variants were examined: octasaccharides and tetrasaccharides with reducing ends of GlcUA (U-HA8 and U-HA4) or GlcNAc (N-HA8 and N-HA4). Among these, HA8 spanned the TM tunnel to elucidate the allosteric correlation between the TM outlet and the GT pocket (Extended Data Fig. 4). Equilibrium conformations were obtained through 800∼900 ns of conventional MD (cMD) or 200 ns of simulated-annealing-accelerated MD (SA-MD) (Supplementary Fig. 5D). The dynamic cross-correlation matrix (DCCM) confirmed the correlations between the TM and GT domains (Extended Data Fig. 5A). However, binding of various HAs did not significantly alter the correlation pattern or GT pocket conformations (Extended Data Fig. 5A-B). Subsequent docking of substrates also revealed no significant differences in binding positions or energies between UDP-GlcUA and UDP-GlcNAc (Extended Data Fig. 5C-D, Extended Data Fig. 6). Therefore, the static conformation of active pocket, even reshaped upon HA binding, still failed to confer substrate specificity, promoting investigation into the enzyme dynamics upon substrate binding.

All-atom MD simulations were performed to compare the binding energies of different substrates to HAS-HA complexes. Given the challenges of achieving equilibrium in systems with membrane proteins and flexible glycans (Supplementary Fig. 5D), we employed a multiple simulated-annealing-MD (MSA-MD) strategy^25, 26^ for conformational sampling, which complements to cMD and SA-MD simulations. Equilibrated conformations with reactive atoms within 2 nm were considered as catalysis-relevant outcomes. After binding with U-HA8 and N-HA4, only alternating substrates could approach HA with low binding energy (Fig. 5A). The HA terminals played a crucial role in regulating specificity which adopted two distinct positions: U-HA4 and N-HA8 fully enter the TM tunnels, allowing for substrate binding at the active sites, while U-HA8 and N-HA4 linger within the active pockets, with substrates binding in the substrate tunnels (Extended Data Fig. 7). Although the two substrates displayed similar affinities at the active sites, strictly alternating specificity was observed within the substrate tunnels (Fig. 5A).

**Fig. 5.**
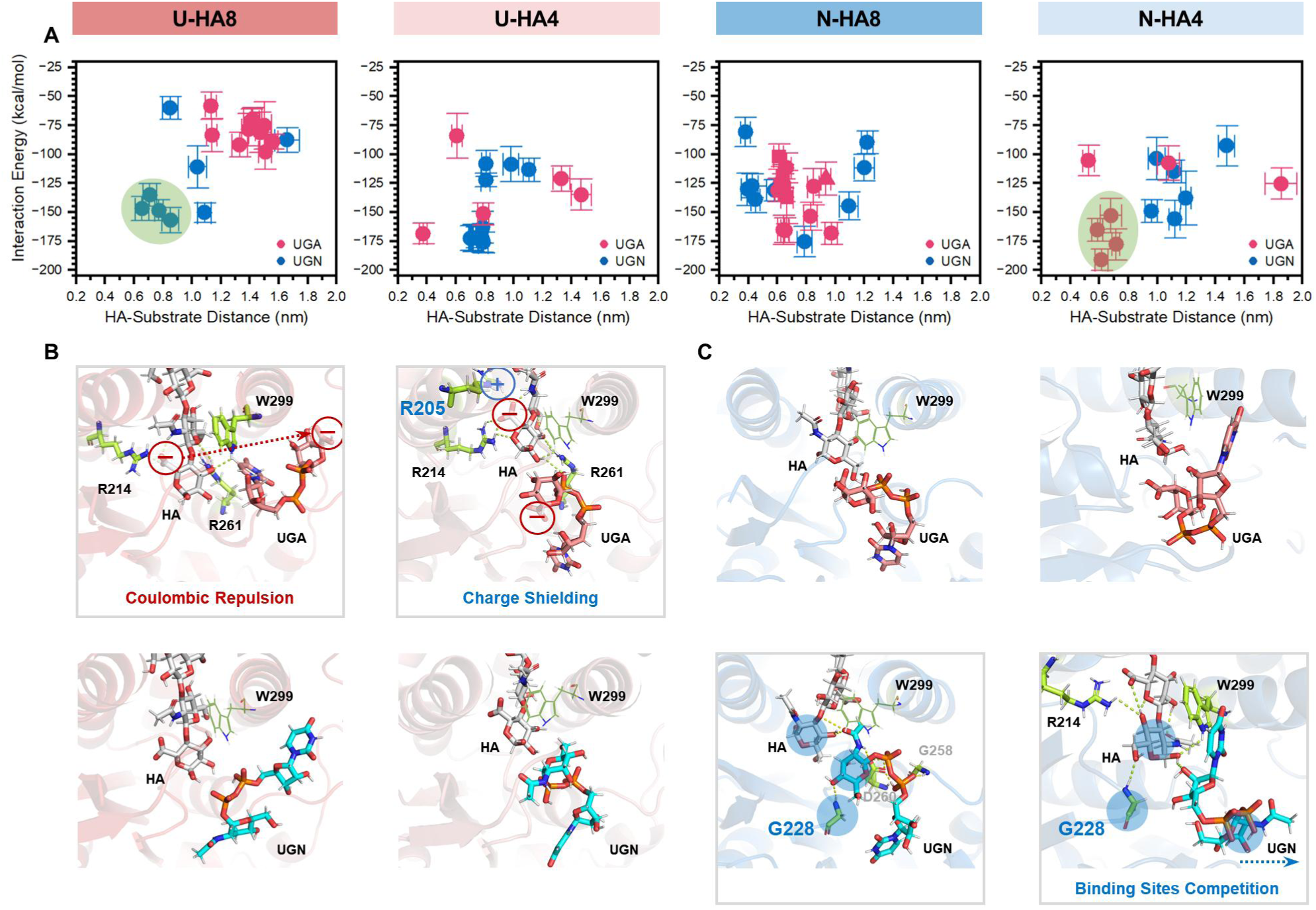
HA terminals regulate the alternating substrate specificity in substrate tunnels. (A) The binding energy between substrate and HAS-HA versus distance between reactive atoms of HA and substrate. (B) Carboxyl group at the HA terminal repulsed UGA through electrostatic forces. (C) GlcNAc at HA terminal competed for UGN binding sites.

When U-HA lingered in the active pocket (U-HA8), its terminal carboxyl group exerted significant Coulombic repulsion on the carboxyl group of the UGA substrate. This trapped the non-alternating substrate in a potential well far from HA (Extended Data Fig. 8), effectively preventing the sugar ring of UGA from accessing the active pocket (Fig. 5B). However, when HA fully entered the TM tunnel (U-HA4), the terminal carboxyl group was shielded by R214 and additional R205, allowing both substrates entry into the active site (Extended Data Fig. 7, Fig. 5B). At this point, although the HA terminal participated in substrate positioning through direct hydrogen bonding, the binding sites and energies for different substrates became less distinguishable (Extended Data Fig. 7, Fig. 5A).

Similarly, when N-HA lingered in the active pocket (N-HA4), the terminal GlcNAc competed with the substrate UGN for binding positions and the binding site G228. This prevents the non-alternating substrate from accessing the active pocket, as suggested by the cryo-EM structure^8^. However, when HA fully entered the TM tunnel (N-HA8), the HA terminal positioned different substrates at similar binding sites through hydrogen bonding, eliminating alternating specificity (Extended Data Fig. 7, Fig. 5C). We thus conclude that the HA terminal lingering in the active pocket consecutively chooses the alternating substrate through Coulombic repulsion and the competition for binding sites, which results in the switching of substrate specificity in HAS catalysis *in vitro*.

### Dynamic C-terminal loop shortened the transport distance for the alternating substrate

We further revealed that the dynamic C-terminal loop (T404∼L417, denoted as C-loop) at the substrate tunnel entrance regulated alternating selectivity by shorten the transport distance of the alternating substrate once HA fully entered TM tunnel. SA-MD simulations indicated that the motions of C-terminal loop facilitated substrate capture and retention (Fig. 6A, Supplementary Material 1). Before substrate binding, the C-loop underwent significant oscillations to capture substrates from solution. After pulling substrate into the tunnel entrance, the C-loop returned to its open conformation. For substrates attempting to escape the active pocket, the C-loop swung to close the substrate tunnel and retain the substrates. Conversely, we infer that such loop dynamics are restricted during regular substrate transport and binding, preventing overcrowding of the active pocket by multiple substrates. Therefore, the transport pathway can be analyzed using the static HAS structure with substrate bound at the active site.

**Fig. 6.**
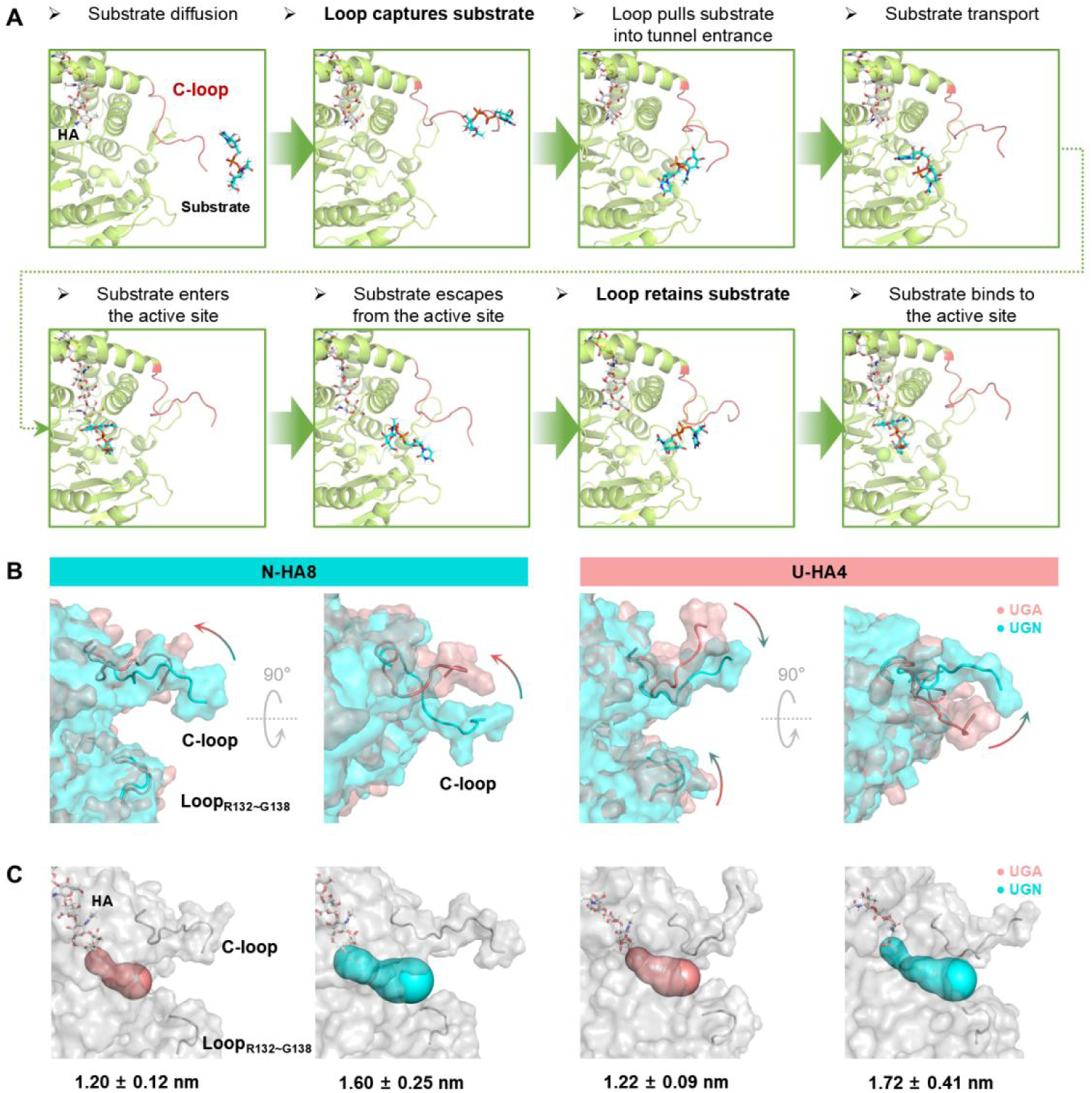
C-terminal loops facilitate alternating substrate specificity in the substrate tunnels. (A) C-loop dynamics regulates substrate capture and retention. (B) Motions of C-loop and loop_R132∼D138_, and (C) length of substrate tunnels upon entry of different substrates. The tunnel starting points are defined by the O atoms of terminal HA sugar ring.

The HAS-HA-substrate complexes with binding energies below −130 kcal/mol and HA-substrate distances less than 1 nm were selected for tunnel analysis using Caver^18^. In HAS-(N-HA8), the C-terminal loop swung upward and laterally away from the substrate tunnel upon alternating substrate binding (Fig. 6B), bringing the active site closer to the protein surface and thus reduces the transport distance (Fig. 6C, Extended Data Fig. 10A). Moreover, both substrates exhibited similar energy trajectories during transport toward the active sites (Extended Data Fig. 10B). Thus, it’s inferred that the alternating substrate UGN has a faster binding rate than UUA. In contrast, for HAS-(U-HA4), upon alternating substrate binding, the C-loop swung not only laterally but also downward from the substrate tunnel, accompanied by the upward movement of loop_R132∼G138_ (Fig. 6B), leading to a longer transport distance for UGN (Fig. 6C). The extended transport pathway may facilitate substrate activation toward its transition state, thereby lowering the activation energy in the intrinsic reaction, which will be further investigated by QM/MM. The C-terminal loop of SeHAS has been reported to significantly influence enzyme activity and HA molecular weight, as demonstrated by truncation and site-directed mutagenesis^27^. As the conserved G138 is presumed to regulate substrate binding^28^, it is reasonable that the coordinated motions of the C-loop and loop_R132∼G138_ modulate the pathway for substrate transport. In conclusion, although the two substrates exhibit similar binding energies once HA fully enters TM tunnel, the dynamic C-terminal loop ensures faster binding kinetics for the alternating substrate in HAS-(N-HA).

## Discussion

Fig. 7 summarizes the process by which SeHAS switches substrate specificity within a single active pocket. The substrate tunnel exhibits a dual mechanism of substrate selection, depending on the positions of HA terminals. Before translocation into the TM tunnel, HA terminals regulate lower binding energies for alternating substrates entering the tunnel. After HA translocation into the TM tunnel, the C-loop forms a shorter transport pathway for the alternating substrate, thereby accelerating its binding kinetics. Once HA fully enters the TM tunnel and the substrate binds to the active site, the hydroxyl group of substrate nucleophilically attacks the C1 of HA terminal, extending the HA chain. SeHAS extends the reducing end of HA, leaving a UDP attached to the nascent HA terminal^12^. This UDP is expected to dissociate and release from the sole UDP-binding region before the entry of UDP-sugar substrate. MD simulations revealed the proximity of HA terminal C1 to the catalytic base D260 (Extended Data Fig. 9), supporting UDP cleavage within the active pocket and justifying the use of HA rather than UDP-HA in our studies of substrate selectivity. The above results present the first molecular insights into the sequential substrate selection within a single active pocket of an enzyme in template-independent polymer synthesis.

**Fig. 7.**
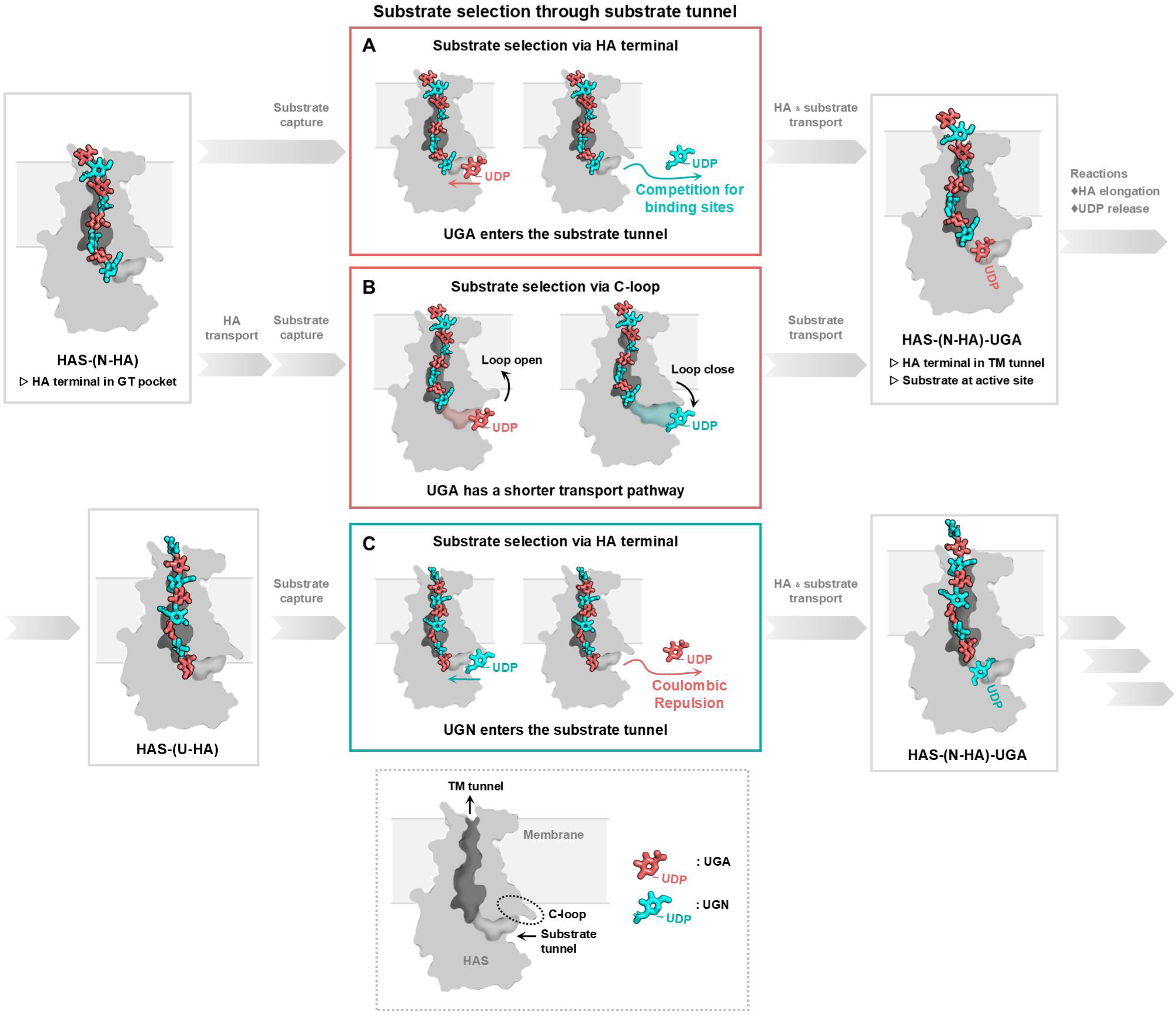
Mechanism of alternating substrate specificity. (A) Lingering N-HA terminal within the active pocket competes for UGN binding sites and allows the entrance of UGA into the substrate tunnel. (B) The C-loop of HAS modulates a shorter pathway for UGA transport when HA fully enters the TM tunnel. (C) Lingering U-HA terminal within the active pocket exerts electrostatic repulsive force on UGA and allows the entrance of UGN into the substrate tunnel.

Above results show that both the active site and the distal regions affect HAS’s substrate specificity, in addition to direct HA-substrate interactions. These distal residues in TM and substrate tunnels are presumed to be common determinants of specificity for the large glycosyltransferase family GT2 in CAZy database (http://www.cazy.org/)^29^. Since the processive synthesis of polysaccharides such as cellulose, chitin, alginate, and poly-NAG also requires continuous polymer transport through the TM tunnel^30^, similar to HA, the TM tunnel may exert allosteric regulation on the active pocket, as observed in HAS. A substrate tunnel connecting the buried active site to the bulk solvent is frequently observed among GT2 members, such as bacterial cellulose synthase BcsA^31^, chitin synthase CaChs2^32^, and mannosyltransferase PcManGT^33^. In BcsA, the dynamic gating loop regulates the opening and closing of the substrate tunnel^31^. In SeHAS, we reveal that dynamic loops in the substrate tunnel regulate transport distance and thereby substrate selection. These findings may inspire further exploration of how substrate tunnel properties influence substrate selectivity in GT2 glycosyltransferases. The complex distal enzyme dynamics provides conceivable mechanisms that determine the well-defined sequence structures of polysaccharides, enhancing our understanding of their critical physiological roles.

## Methods

### Plasmids and strains

Plasmid pMBAD-*sse*AB containing *hasA* gene of *Streptococcus equisimilis* was kindly provided by Prof. Yu Huimin, Tsinghua University, China^34^. A 6×histidine tag was inserted at the N-terminal of HAS by GENEWIZ (Suzhou, China). The site-directed mutagenesis library was constructed using Gibson Assembly^35^. All primers were listed in Supplementary Table 3. Library fragments were amplified by PCR using Phanta Max Super-Fidelity DNA Polymerase from Vazyme (Nanjing, China). The vectors were digested with NcoI and XbaI for mutagenesis targeting residues 34 to 345, and with BamHI and SfiI for residues 6 to 17. Restriction endonucleases and Gibson Assembly^®^ Master Mix were from NEB (Ipswich, USA). All plasmids were transformed into *E.coli* DH5α (Vazyme, Nanjing, China).

### *In vivo* synthesis of HA

Ampicillin selection agar medium: Luria-Bertani (LB) medium (10 g/L tryptone, 5 g/L yeast extract, 10 g/L NaCl) containing 15 g/L agar and 0.2 g/L ampicillin; Fermentation medium: LB medium containing 0.2 g/L ampicillin, 20 mmol/L MgCl_2_.

Recombinant *E.coli* DH5α was selected after incubation at 37℃ for 16 hours on the ampicillin selection agar plate. The selected colony was inoculated into 3 mL of fermentation medium and incubated for 12 hours at 37℃, 200 rpm. Subsequently, 0.5 mL of the culture was transferred into 50 mL fermentation medium in 500 mL shake flasks and incubated at 37℃, 200 rpm. HAS expression was induced with 0.1 g/L arabinose when OD_600_ reached 0.8. Cultivation continued for an additional 5 hours before adding 10 g/L glucose to enhance HA synthesis, and HA was harvested after another 16 hours.

### Preparation of HAS-embedded liposome

*E.coli* DH5α was harvested 5 hours after induction (OD_600_=1.2∼1.5) by centrifugation at 12,000×g for 10 minutes at 4℃. Collected cells were washed twice with PBS-glycerol buffer (pH 7.2, 1.8 mmol/L KH_2_PO_4_, 8 mmol/L Na_2_HPO_4_, 137 mmol/L NaCl, 2.7 mmol/L KCl, 10% glycerol), and resuspended to 1/10 of the original fermentation volume. The suspension was supplemented with 5 mmol/L DTT, 1 mmol/L PMSF, 1× protease inhibitor cocktail (Beyotime, Shanghai, China), and then sonicated at 200 W (3 s on, 3 s off) for 20 minutes in an ice bath for cell disruption. The lysate was centrifuged at 12,000×g for 30 minutes at 4℃ to remove the cell debris. The supernatant was then supplemented with 1 mmol/L PMSF, 1× protease inhibitor cocktail, and 1 mmol/L DTT, followed by ultracentrifugation (Optima XPN-100, Beckman) at 200,000×g for 2 hours at 4℃ to collect the membrane fragments. The pale-yellow lipid film obtained was washed twice with PBS-glycerol buffer, resuspended to 1/20 of the original fermentation volume, supplemented with 1 mmol/L DTT, and stored at −80℃.

### TEM and DLS characterization of liposome

The morphology of liposomes was characterized by negative staining transmission electron microscopy (TEM). The liposomes were diluted with 10% glycerol to a total protein concentration of 0.4∼1 g/L, as determined by the BCA protein assay kit (Solarbio, Beijing, China). 6 μL of diluted sample was applied to a 230-mesh standard carbon-coated grid. Excess liquid was absorbed by filter paper after 3 minutes, and the grid was allowed to air-dry for 2 minutes. Then 6 μL of 2% phosphotungstic acid stain (Solarbio, Beijing, China) was added. Excess liquid was again removed by filter paper after 2 minutes, and the grid was left to air-dry for 12 hours. The TEM (JEM-2010, JEOL) was operated at an acceleration voltage of 200 kV.

The size of liposomes was determined by dynamic light scattering (DLS) with Zetasizer Nano ZS-90 (Malvern). The liposomes were diluted as described above.

### Purification and SDS-PAGE analysis of HAS

The HAS-embedded liposomes with a total protein concentration of 2 g/L were solubilized with 5 mg/mL detergent DDM for 2 hours at 100 rpm, 4℃. The insolubles were removed by ultracentrifugation (Optima MAX-XP, Beckman) for 2 hours at 400,000×g, 4°C. HAS was then purified using the HisTrap HP pre-packed column (Cytiva) on the ÄKTA Pure system, with PBS-glycerol buffer containing 0.5 mg/mL DDM and 1 mmol/L DTT as the equilibration buffer, equilibration buffer containing 30 and 300 mmol/L imidazole as the loading and elution buffer, respectively. Elution was performed isocratically at 4℃ with a flow rate of 1 mL/min. The fractions of purified HAS were subsequently concentrated and desalted by Amicon Ultra Centrifugal Filter 30 kDa MWCO (Milliporea) using the equilibration buffer. The concentrated HAS solution at 1 mg/mL was loaded onto a 12% Tris-glycine gel (Beyotime, Beijing, China) for SDS-PAGE. The protein band at 42 kDa was identified as HAS rather than the theoretical value of 48 kDa^11^.

### *In vitro* synthesis of HA

The reaction mixture containing SeHAS-embedded liposomes with HAS at a final concentration of 0.4 μmol/L, UDP-GlcUA and UDP-GlcNAc substrates, 20mmol/L MgCl_2_ and 1 mmol/L DTT was prepared in PBS-5% glycerol buffer (pH 7.2, 1.8 mmol/L KH_2_PO_4_, 8 mmol/L Na_2_HPO_4_, 137 mmol/L NaCl, 2.7 mmol/L KCl, 5% glycerol). The reaction was conducted at 37℃ with shaking at 200 rpm for 12 hours, and terminated by heating in a 100 ℃ water bath for 5 minutes.

### Purification of HA

The enzymatic reaction mixture or the fermentation culture was centrifuged at 12,000×g for 10 minutes at 4°C, and HA in the supernatant was precipitated by adding twice the volume of anhydrous ethanol. After incubation at 4°C for 2 hours, HA was collected by centrifugation at 12,000×g for 15 minutes at 4°C and redissolved in 0.2 mol/L NaCl. After removing insolubles by centrifugation at 12,000×g for 15 minutes at 4℃, HA was reprecipitated with ethanol to obtain a crude HA product for quantification and gel filtration chromatography (GFC) analysis.

The crude HA was further purified by chromatography on the ÄKTA Pure system for structural characterization. HA was first purified by anion-exchange chromatography using a Hitrap Q FF pre-packed column (Cytiva), with 50 mmol/L Tris-HCl (pH 8.0) from Solarbio (Beijing, China) as equilibration buffer, equilibration buffer containing 0.1 and 1 mol/L NaCl as loading and elution buffer, respectively. Elution was conducted at 4°C using a linear gradient across 10 column volumes, at a flow rate of 5 mL/min. Size exclusion chromatography was then performed using a SUPEROSE 6 Increase 10/300 GL column (Cytiva), with ultrapure water as the mobile phase at a flow rate of 0.5 mL/min. Absorbances at 210 nm and 280 nm were simultaneously monitored, and the fractions with high absorption at 210 nm and low absorption at 280 nm were identified as HA.

### Quantification, GFC analysis and HAase-mediated digestion of HA

The crude or purified HA was quantified by CTAB turbidimetric method^36^.

Molecular weight of HA was characterized by GFC analysis using the PL aquagel-OH columns (Agilent), monitored by photodiode array (PDA) or refractive index (RI) detector. The triple tandem column consisted of PL aquagel-OH 20, PL aquagel-OH 40 and PL aquagel-OH MIXED-M. The mobile phase was 0.2 mol/L NaCl at a flow rate of 1 mL/min.

HA was specifically digested by bovine testicular hyaluronidase (HAase) from Sigma Aldrich (St. Louis, USA). 1 mg/mL of HA was degraded by 10 U/mL of HAase for 12 hours at 37℃ in ultrapure water or 0.2 mol/L NaCl.

### Structural characterization of HA

The consumption of UDP-GlcUA and UDP-GlcNAc substrates, as well as the production of GlcUA and GlcNAc monosaccharides from substrate hydrolysis were determined by anion-exchange high-performance liquid chromatography (AEX HPLC) using the NH2P-50 4E column (Shedox). The mobile phase was PBS buffer (pH 7.2, 1.8 mmol/L KH_2_PO_4_, 8 mmol/L Na_2_HPO_4_, 137 mmol/L NaCl, 2.7 mmol/L KCl) at a flow rate of 1 mL/min. PDA detector monitored the absorbances at 280nm, 260 nm and 205 nm to detect proteins, substrates, monosaccharides and HA, respectively.

The monomeric composition of HA was determined by proton nuclear magnetic resonance (^1^H NMR) spectrum using 600 MHz spectrometer JNM-ECA60 (JEOL). The high-molecular-weight HA was dissolved in D_2_O to 1∼2 mg/mL, and was tested immediately after the addition of 75 mmol/L NaOD at room temperature. The offset frequency, sweep width, relaxation delay were set to 3.0 ppm, 3.2 ppm, 20 s, respectively, and 32∼64 scans were acquired.

The sequence of HA was determined by enzymatic digestion using chondroitinase ABC (ChABC) from *Proteus vulgaris* (Sigma Aldrich, St. Louis, USA) and subsequent electrospray ionization mass spectrometry (ESI-MS) analysis using LCMS-8040 (Shimadzu). The ChABC was dissolved in 50 mmol/L acetate (pH 7.0) to 100 U/L, and 0.02% of bovine serum albumin (BSA) was added. The potassium phosphate buffer salts and stabilizers were removed using Ultra Centrifugal Filter 30 kDa MWCO. 1 mg/mL of HA was degraded by 100 U/L of ChABC for 3 hours at 37℃. After confirming full digestion by GFC, the ChABC was removed by Ultra Centrifugal Filter 10 kDa MWCO. The degradation products were analyzed by ESI-MS and further ESI-MS/MS at negative mode, with 95% acetonitrile at 0.4 mL/min as mobile phase. The fragment ions produced by ESI-MS/MS were predicted by CFM-ID 3.0^37^.

### Structure prediction, molecular docking, tunnel prediction and allosteric analysis of HAS

The 3D structure of SeHAS monomer and dimer were predicted by AlphaFold2^13, 16^ on the ColabFold^38^ platform. Protein-protein docking was performed by HADDOCK^14^. The orientations of SeHAS in membranes were predicted by PPM 3.0^15^. Molecular docking was performed using Autodock Vina^39^. The structures of HA were modeled by CHARMM-GUI Glycan Modeler^40^, and structures of substrates were from PubChem database^41^. Bindging sites for Mg^2+^ were predicted by MIB2^42^. Tunnel prediction was performed via Caver Web^18^ or Caver PyMOL plugin (version 3.0.3)^43^. Tunnel analysis of the AlphaFold-predicted structures used default parameters, while the MD-equilibrated structures used the following parameters: minimum probe radius = 0.14 nm, shell radius = 0.55 nm, and shell depth = 0.35 nm. The allosteric analysis was conducted on Ohm web server^21^.

### MD simulations

All molecular dynamics (MD) simulations were conducted using Gromacs-2019.4, with CHARMM36m^44^ forcefield for all-atom simulation and Martini^45^ forcefield for coarse-grained simulation. Parametrization of substrates and HA was performed by FFParam^46^ and CHARMM-GUI Glycan Modeler^40^, respectively. The input files for simulations were constructed using CHARMM-GUI^47^. HAS was embedded into lipid bilayers (Supplementary Table 1-2) and solvated with TIP3P water, supplemented with 0.15 mol/L NaCl, in a simulation box of 12 nm × 12 nm × 12.5 nm.

Energy minimization was executed using steepest descent algorithm. And 6 iterations of equilibration were conducted under the NPT ensemble with restraints on proteins, lipids, HA and substrates gradually decreased, a procedure instructed by standard CHARMM-GUI-derived input files. The production simulations were carried out under the NPT ensemble, with temperature controlled at 303.15 K and pressure controlled at 1 bar using V-rescale and Parrinello-Rahman method, respectively. Semi-isotropic pressure coupling was employed to make the pressure in the membrane plane independent from the z-axis. Periodic boundary conditions were applied in the x, y, and z directions. The simulation time step was set to 20 fs for coarse-grained MD and 2 fs for all-atom MD.

In simulated-annealing-accelerated MD (SA-MD), the production simulation initiated with a simulated annealing (SA) procedure to search for the globally optimal conformation and thereby facilitate the equilibration. Different SA procedures were illustrated in Supplementary Fig. 4, with SA_single procedure employed for SA-MD simulations. In multiple simulated-annealing-MD (MSA-MD), parallel runs using SA_single procedure were performed, with complementary SA_Twice and SA_Long procedure, to obtain more than 8 catalysis-relevant conformations for each HAS-HA-substrate complex. Catalysis-relevant conformations were defined as those at equilibrium, where the distance between the reactive atoms of HA (C1) and substrates (hydroxyl O on C3 and C4 for UGN and UGA, respectively) was less than 2 nm. The total simulation time for MSA-MD using SA_single, SA_Twice, SA_Long procedure was 35 ns, 50 ns and 35 ns, respectively.

The analysis of trajectory was performed using MDAnalysis^48^.

## Supporting information

Animations of C-loop dynamics

## Acknowledgment

This work was supported by National Natural Science of China under grant no.22120102003 and the National Key Research and Development Program of China (no. 2021YFC2102801).

## Extended data

**Extended Data Fig. 1.**
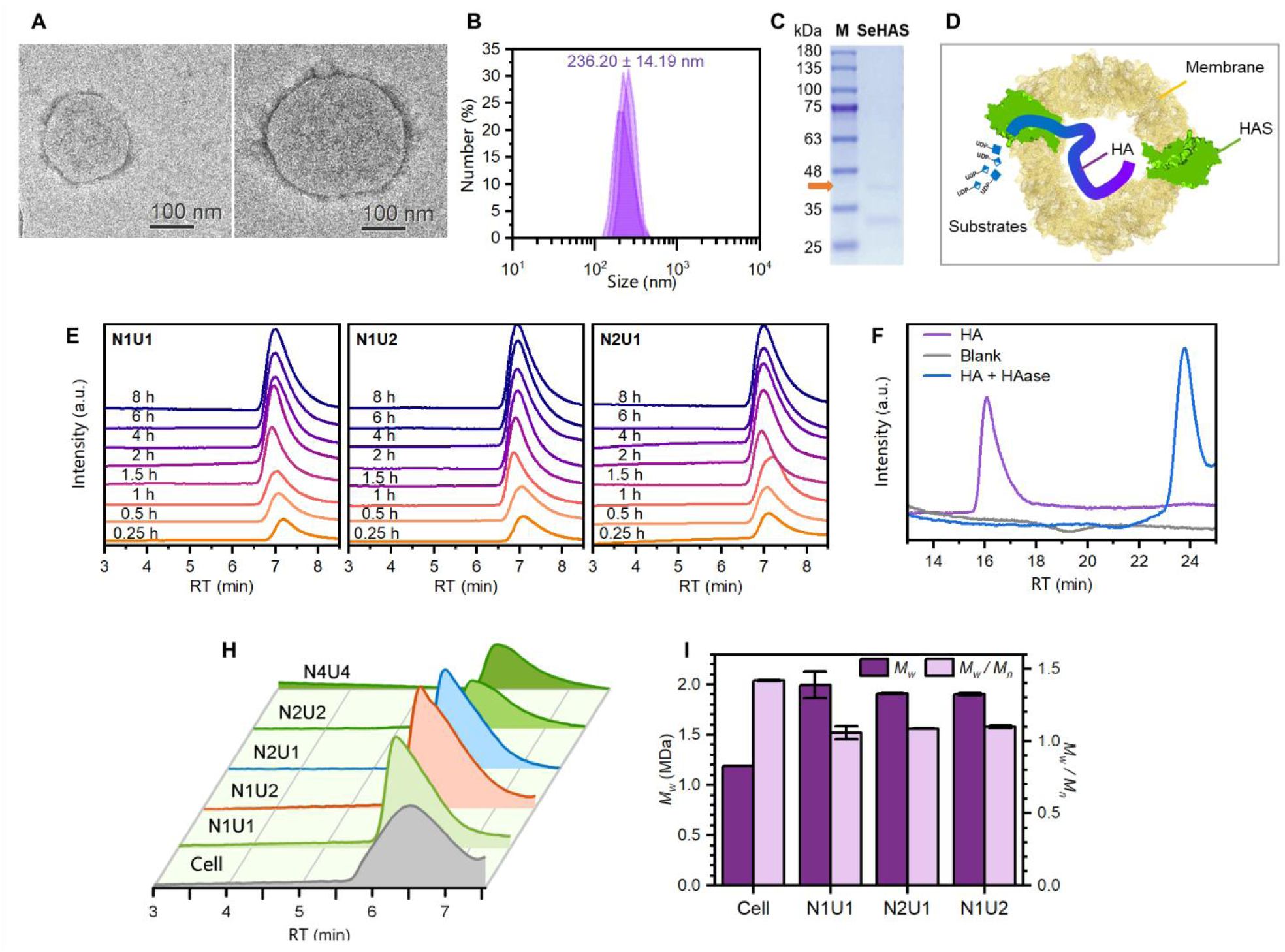
Characterization of HAS catalyst and catalysis *in vitro.* (A) TEM images, (B) DLS analysis, (C) SDS-PAGE, and (D) catalysis processs of HAS-embeded liposomes. (E) GFC profiles over reaction time. (F) HA product was degraded by HAase, with N1U1 as an example. Blank: reaction mixture without substrate. (H) GFC profiles, and (I) *M_w_* and *M_w_/M_n_* at different substrate concentrations and ratios. “N”: UGN, “U”: UGA, numbers indicate original concentrations (mmol/L). Column for (E) and (H) was PL aquagel-OH MIXED-H, and for (F) is a triple tandem of PL aquagel-OH columns.

**Extended Data Fig. 2.**
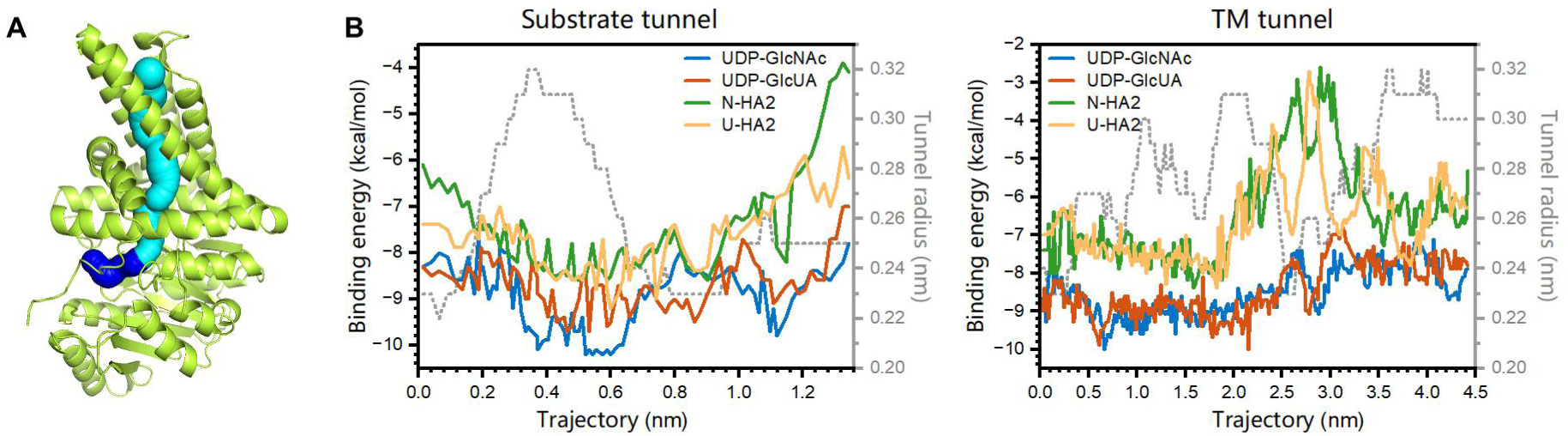
(A) Tunnels in the SeHAS-Mg^2+^ complex, SeHAS structure predicted by AlphaFold2. Blue: substrate tunnel, cyan: TM tunnel. (B) Transport of ligands in tunnels. N-HA2 and U-HA2 represent HA disaccharide with terminal GlcNAc and GlcUA respectively.

**Extended Data Fig. 3.**
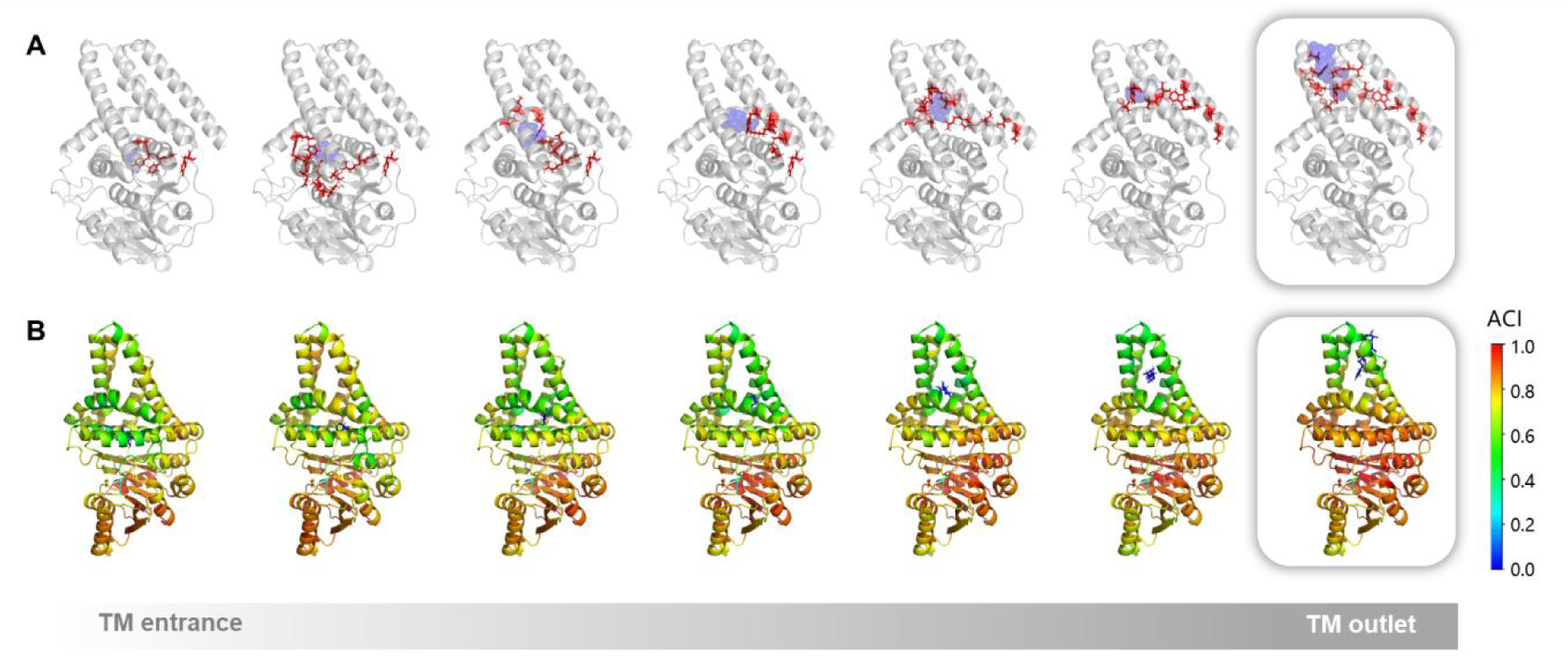
Allosteric correlations between the GT pocket and each of the seven glycosyl binding sites along the TM tunnel. (A) Glycosyl (blue spheres) and the allosteric pathway (red sticks). (B) Allosteric coupling intensity (ACI) of HAS residues.

**Extended Data Fig. 4.**
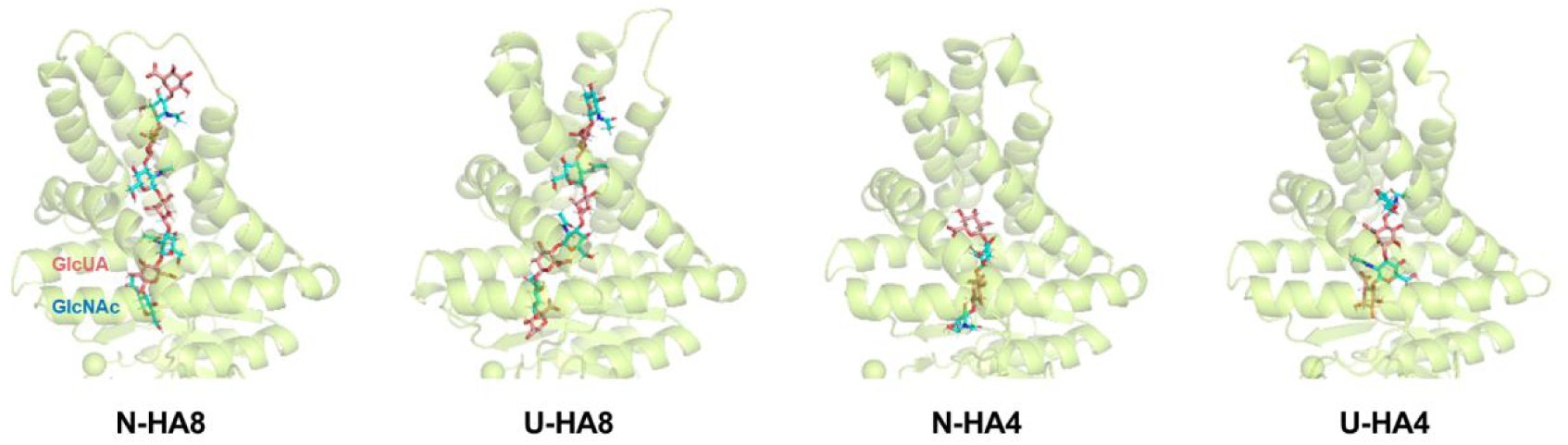
Four HA variants for MD simulations and their equilibrium conformations. Pink and cyan glycosyl represent the GlcUA and GlcNAc unit respectively.

**Extended Data Fig. 5.**
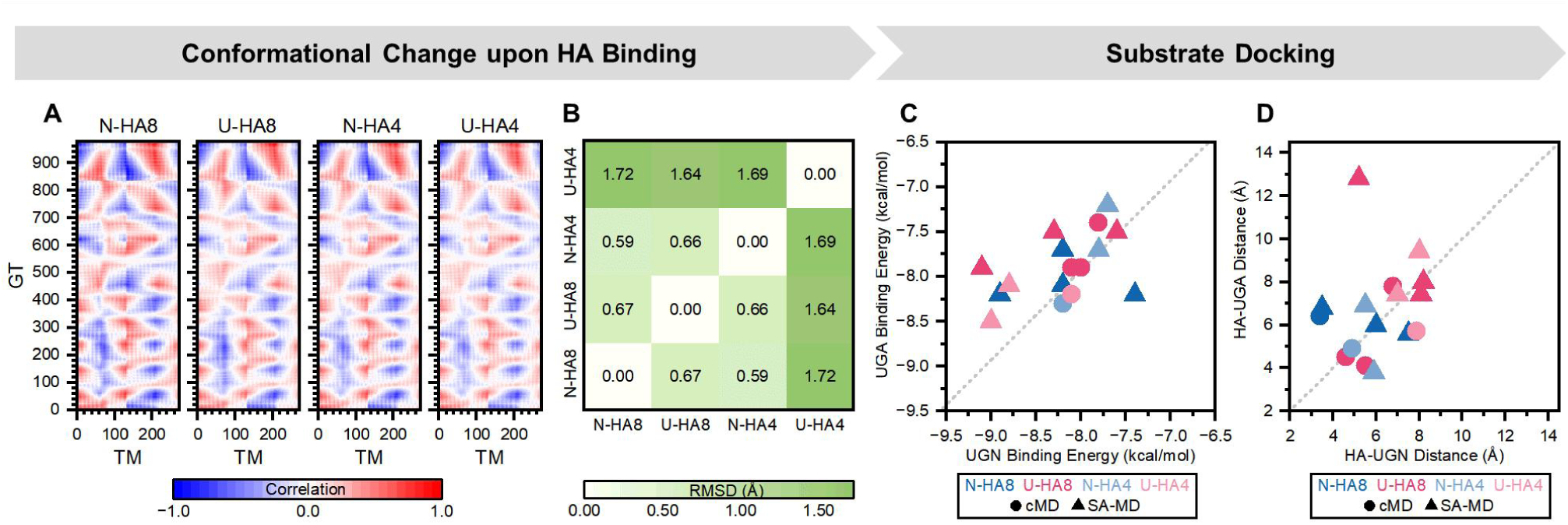
Indistinguishable substrate affinity in static active pocket of HAS-HA complex. (A) DCCM of GT and TM atoms. (B) RMSD of GT domains. (C) The binding energies of different substrates. (D) The distances between reactive atoms of HA and different substrates. Grey dash lines correspond to *y = x*.

**Extended Data Fig. 6.**
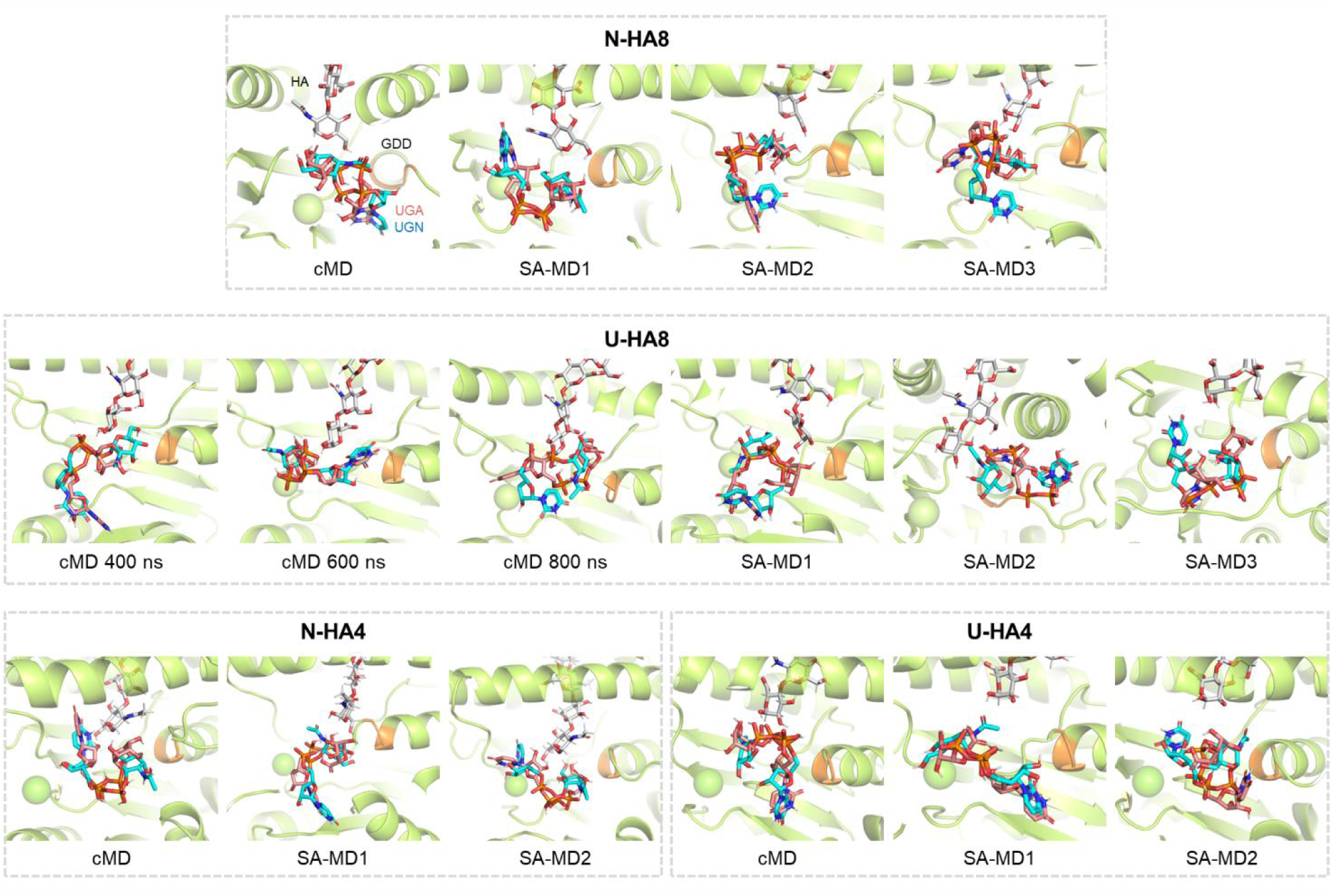
Different substrates docked within HAS-HA complexes shared binding positions. Cyan: UGN, pink: UGA, grey: HA, orange residues: the GDD motif for catalysis.

**Extended Data Fig. 7.**
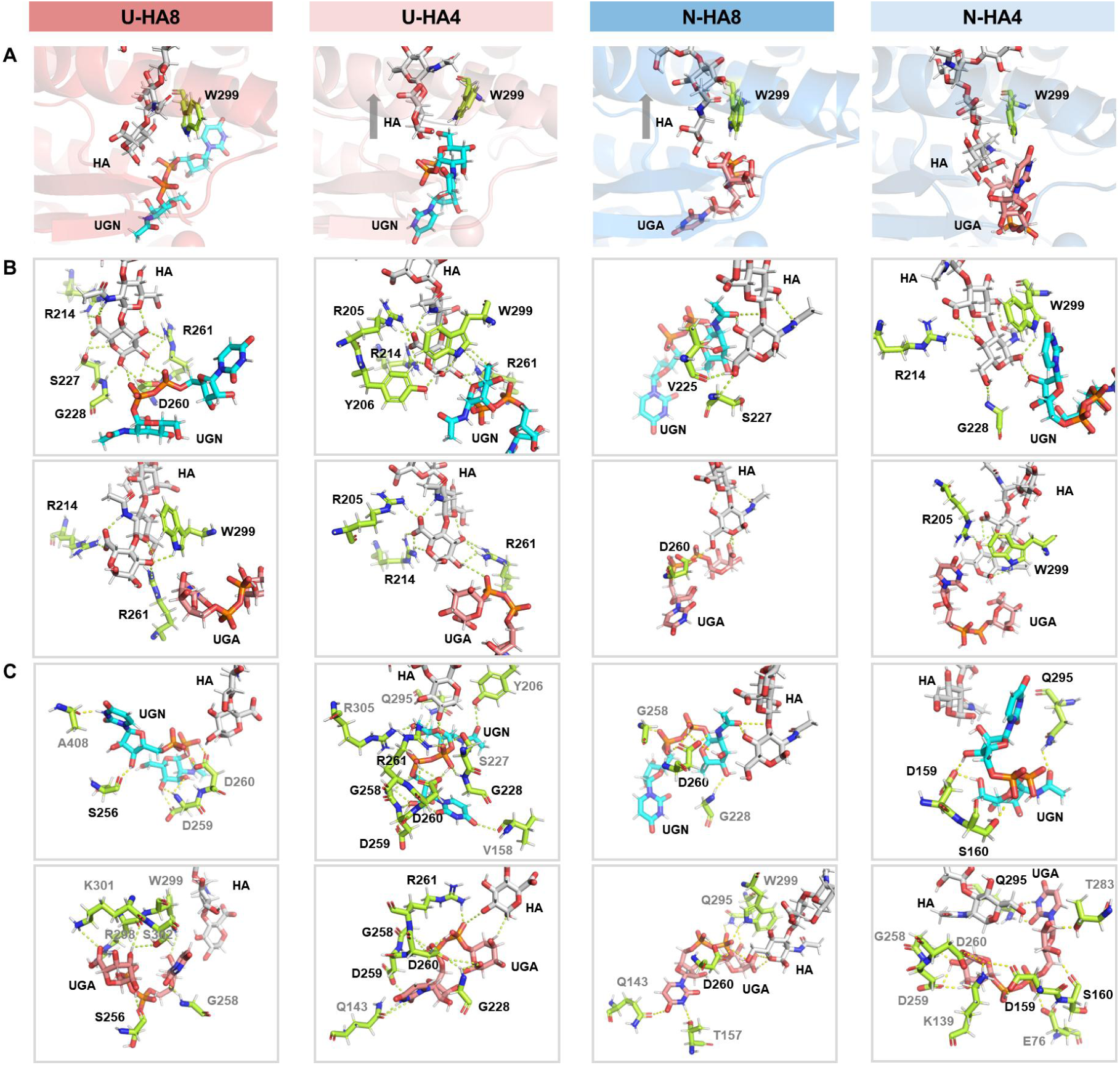
(A) HA terminals within the TM tunnel or at the active pocket. (B) Binding sites for HA terminals. (C) Binding sites of substrates. Black-ticked residues: common binding sites shared by two substrates.

**Extended Data Fig. 8.**
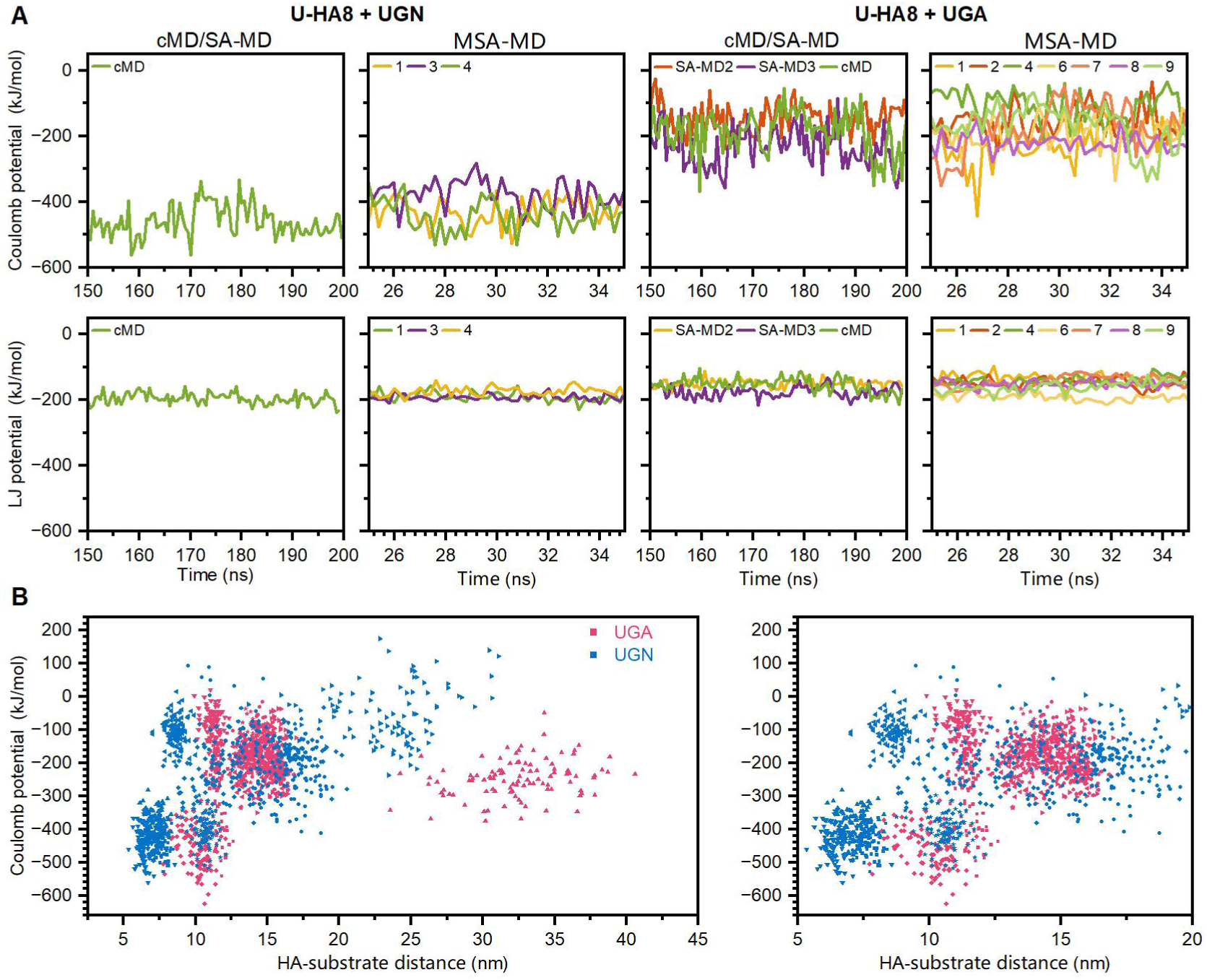
Coulomb potential of different substrates within HAS-(U-HA8)-substrate complexes. (A) Coulomb and LJ potentials at equilibrium. (B) Coulomb potential versus distance between reactive atoms of HA and substrates.

**Extended Data Fig. 9.**
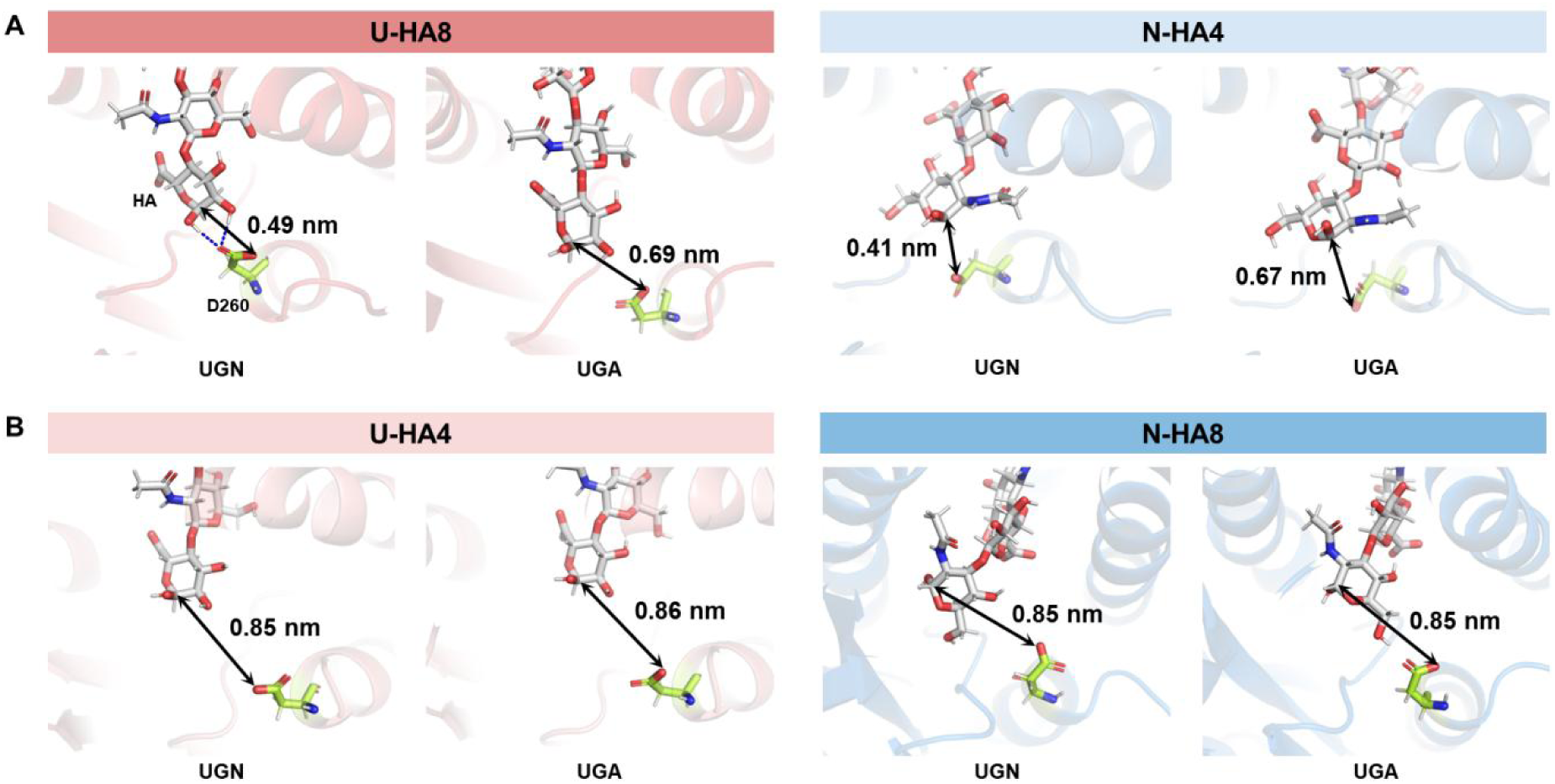
Distance between C1 of HA terminal and hydroxyl O of D260 side chain.

**Extended Data Fig. 10.**
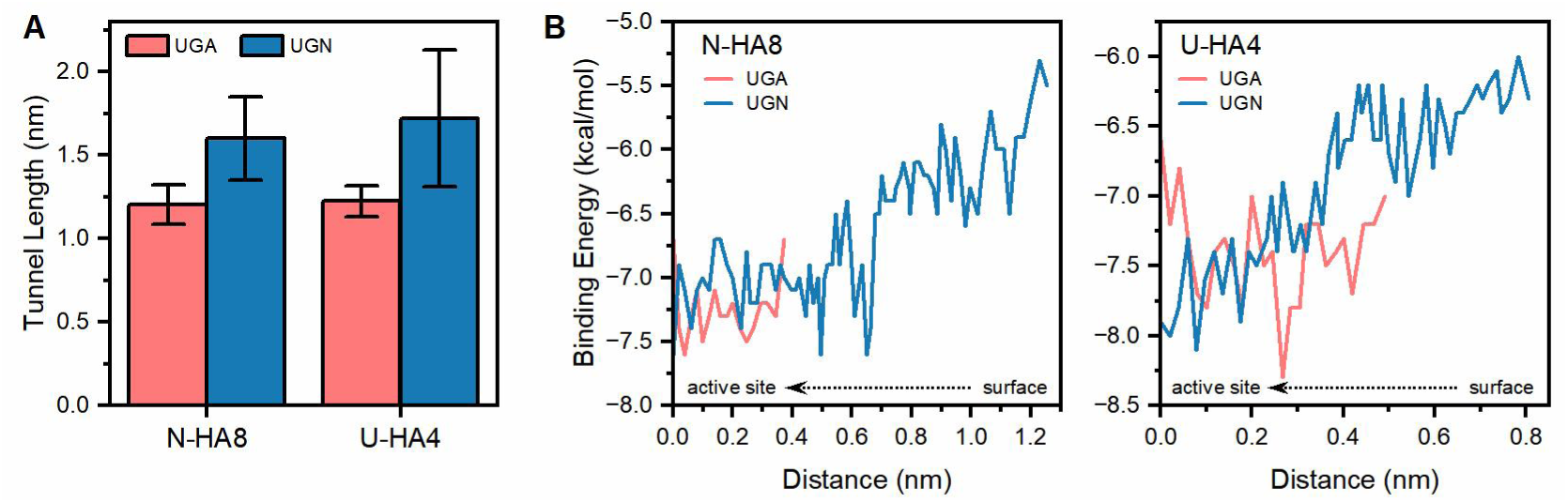
(A) Average length of substrate tunnels and (B) energy trajectory of substrate transport. The tunnel starting points in (A) and (B) are defined by the O atoms of the terminal HA sugar ring and the reactive hydroxyl group in substrate, respectively.

## Notes

### Competing Interest Statement

The authors have declared no competing interest.

