## Supplementary Tables and Figures for "Substrate tunnel regulates substrate specificity switching in hyaluronan synthase"

### Contents

Supplementary Table 1 Compositions of membranes for coarse-grained MD ............ S-1
Supplementary Table 2 Composition of membrane for all-atom MD ............ S-2
Supplementary Table 3 Mutagenesis primers ............ S-2
Supplementary Fig. 1 AEX HPLC analysis ............ S-3
Supplementary Fig. 2 Degradation of HA by chondroitinase ABC. ............ S-4
Supplementary Fig. 3 ESI-MS/MS spectrum ............ S-5
Supplementary Fig. 4 Simulated annealing procedure ............ S-5
Supplementary Fig. 5 Time profiles of MD simulations ............ S-6

Supplementary Table 1 Compositions of membranes for coarse-grained MD

| Membrane | Number of lipids in Upperleaflet | | | Number of lipids in Lowerleaflet | | |
|---|---|---|---|---|---|---|
| | CDL2 | POPG | POPE | CDL2 | POPG | POPE |
| CL | 99 | 0 | 36 | 88 | 0 | 32 |
| IM | 21 | 42 | 147 | 20 | 40 | 140 |
| PE/PG | 0 | 72 | 168 | 0 | 66 | 154 |
| PE | 0 | 0 | 235 | 0 | 0 | 216 |

Supplementary Table 2 Composition of membrane for all-atom MD

| **Layer** | **Number of lipids** | | | | |
|---|---|---|---|---|---|
| | **PMCL2** | **TYCL2** | **DYPE** | **DPPE** | **PYPE** |
| **Upperleaflet** | **72** | 27 | 18 | 9 | 9 |
| **Lowerleaflet** | 64 | 24 | 16 | 8 | 8 |

Supplementary Table 3 Mutagenesis primers

| **Primer** | | **Sequence (5’ → 3’)** |
|---|---|---|
| Cut1[a] | Fwd | TAAGAAGGAGATATACATACCCATGGGTATGCGTACCCTGAAGAAC |
| | Rev | TTAGCATCCTGATAAGAGCTCTAGATTACAGCAGTTTTTTACGCG |
| Cut2 | Fwd | GCATTTTTATCCATAAGATTAGCGGATCCTACCTGACGCTTTTTATCG |
| | Rev | AATCGCGGCCACTTTATACTGGCCTGCACGGCCTTTAAACGG |
| N6A | Fwd | ACCCTGAAGGCTCTGATTACGGTGGTTGCATTTAGCATTTTTTGGGTT |
| | Rev | CACCGTAATCAGAGCCTTCAGGGTACGCATACCCAT |
| V10A | Fwd | ATTACGGCTGTTGCATTTAGCATTTTTTGGGTTCTGCTGATTTATGTG |
| | Rev | AATGCTAAATGCAACAGCCGTAATCAGGTTCTTCAGGGTACG |
| F13A | Fwd | ATTACGGTGGTTGCAGCTAGCATTTTTTGGGTTCTGCTGATTTATGTG |
| | Rev | AATGCTAGCTGCAACCACCGTAATCAGGTTCTTCAGGGTACG |
| W17A | Fwd | ATTTTTGCTGTTCTGCTGATTTATGTGAACGTTTACCTGTTCGGCGCG |
| | Rev | ATAAATCAGCAGAACAGCAAAAATGCTAAATGCAACCACCGTAATCAG |
| L34A | Fwd | AAAGGCAGCGCTAGCATTTATGGTTTTCTGCTGATCGCG |
| | Rev | AAAACCATAAATGCTAGCGCTGCCTTTCGCGCCGAA |
| S35A | Fwd | AAAGGCAGCCTGGCTATTTATGGTTTTCTGCTGATCGCG |
| | Rev | AAAACCATAAATAGCCAGGCTGCCTTTCGCGCCGAA |
| Y37A | Fwd | CTGAGCATTGCTGGTTTTCTGCTGATCGCGTATCTGC |
| | Rev | CAGCAGAAAACCAGCAATGCTCAGGCTGCCTTT |
| G38A | Fwd | CCTGAGCATTTATGCTTTTCTGCTGATCGCGTATCTGC |
| | Rev | CAGCAGAAAAGCATAAATGCTCAGGCTGCCTTT |
| I42A | Fwd | TTTCTGCTGGCTGCGTATCTGCTGGTTAAAATGAGCC |
| | Rev | CAGCAGATACGCAGCCAGCAGAAAACCATAAATGCTC |

| Primer | | Sequence (5' → 3') |
|---|---|---|
| L334A | Fwd | ATGTTCATGATGGCTGTGTACAGCGTGGTTGACTTCTTT |
| | Rev | GCTGTACACAGCCATCATGAACATGCTAACTTCCAGAATGGT |
| S337A | Fwd | CTGGTGTACGCTGTGGTTGACTTCTTTGTGGGCAAC |
| | Rev | GAAGTCAACCACAGCGTACACCAGCATCATGAACATGC |
| V338A | Fwd | CTGGTGTACAGCGCTGTTGACTTCTTTGTGGGCAAC |
| | Rev | GAAGTCAACAGCGCTGTACACCAGCATCATGAACATGC |
| F341A | Fwd | GTGGTTGACGCTTTTGTGGGCAACGTGCGC |
| | Rev | GTTGCCCACAAAAGCGTCAACCACGCTGTACACCAGCAT |
| F342A | Fwd | GTGGTTGACTTCGCTGTGGGCAACGTGCGC |
| | Rev | GTTGCCCACAGCGAAGTCAACCACGCTGTACACCAGCAT |
| N345A | Fwd | TTTGTGGGCGCTGTGCGCGAGTTCGACTGG |
| | Rev | GAACTCGCGCACAGCGCCCACAAAGAAGTCAACCAC |

a: Cut1 and Cut2 are vector primers for mutagenesis targeting residues 34 to 345 and 6 to 17, respectively.

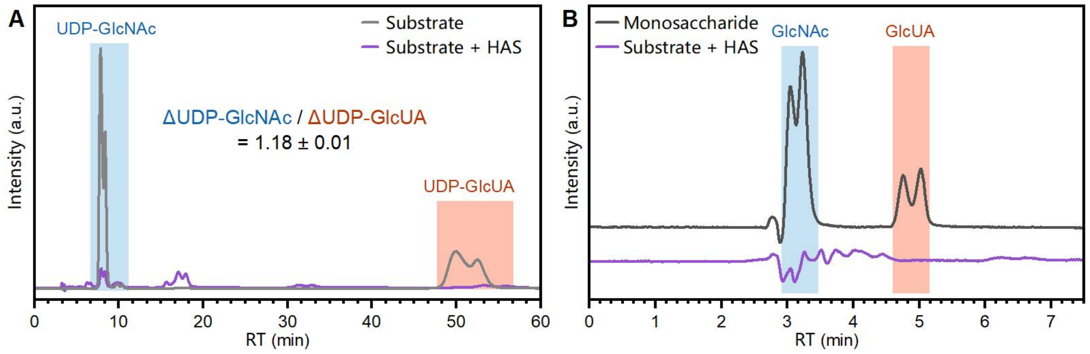


Supplementary Fig. 1 AEX HPLC analysis of (A) the molar ratio of substrate consumption and (B) the hydrolysis of substrates, with N1U1 as an example.

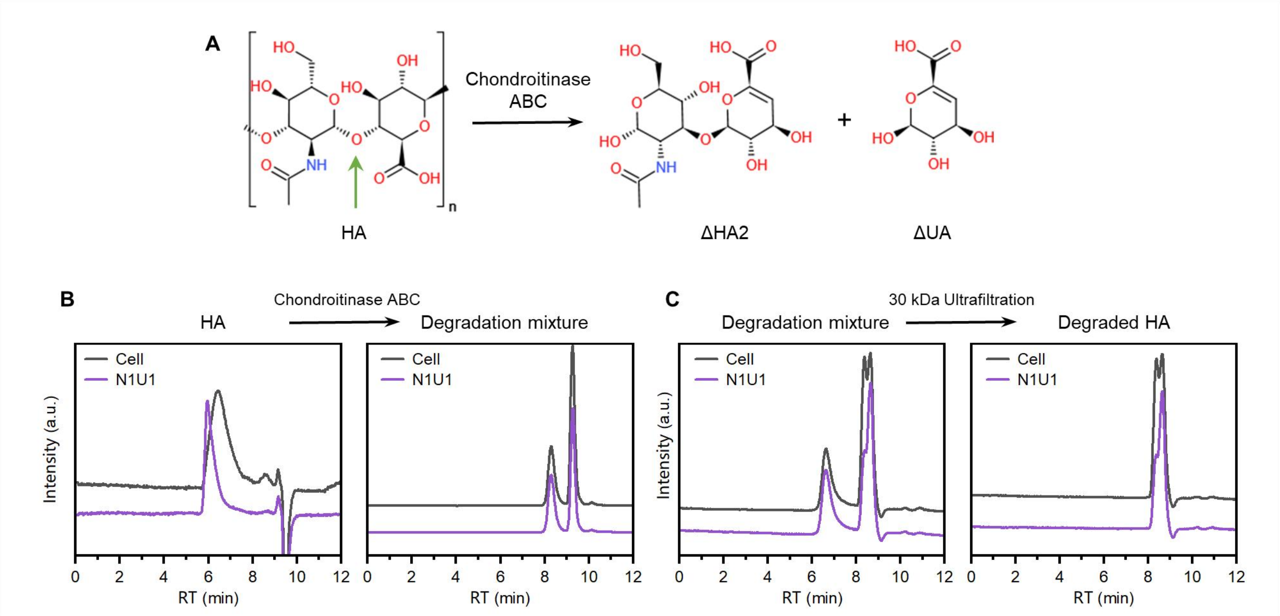


Supplementary Fig. 2 Degradation of HA by chondroitinase ABC. (A) Reaction. GFC analysis determined (B) the degradation of macromolecules and (C) the molecular weight of the degradation products, using columns PL aquagel-OH MIXED-H (6 kDa to 10 MDa) and PL aquagel-OH 30 (100 Da to 60 kDa), respectively.

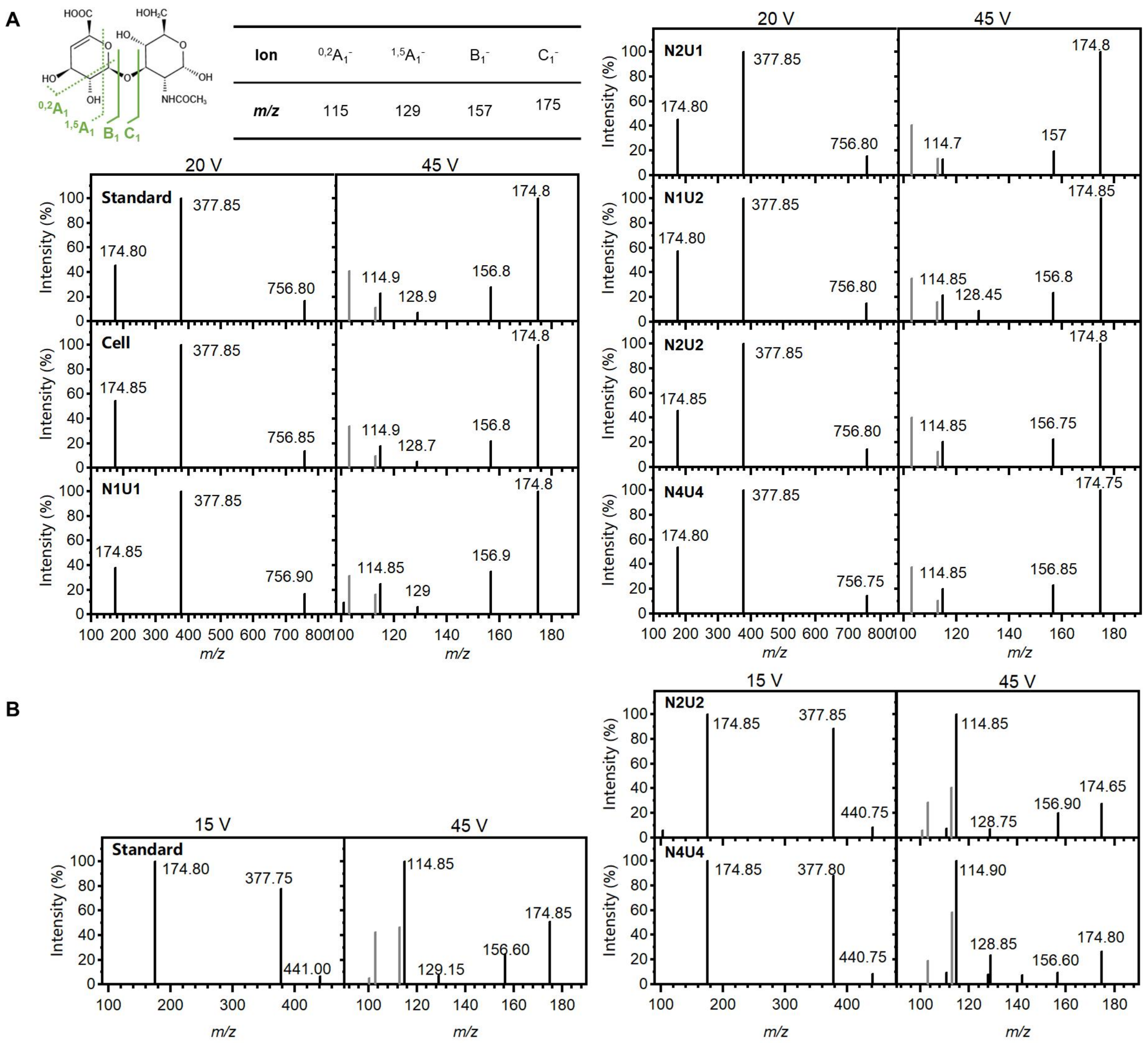


Supplementary Fig. 3 ESI-MS/MS spectrum of the (A) *m/z*=757 ions and (B) *m/z*=441 ions at low and high collision energies.The *m/z*=757 and *m/z*=441 correspnding to [ΔHA2]$_2^-$ and ions of ΔHA2-ΔUA complex. Unlabeled gray peaks are from GlcNAc units. "N": UGN, "U": UGA, numbers indicate original concentrations (mmol/L).

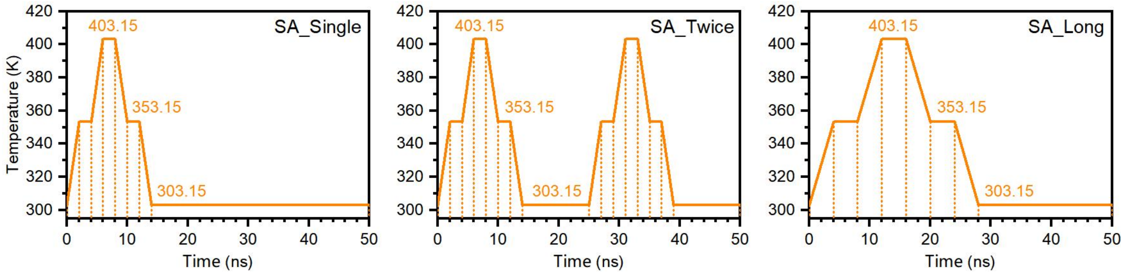


Supplementary Fig. 4 Simulated annealing procedure.

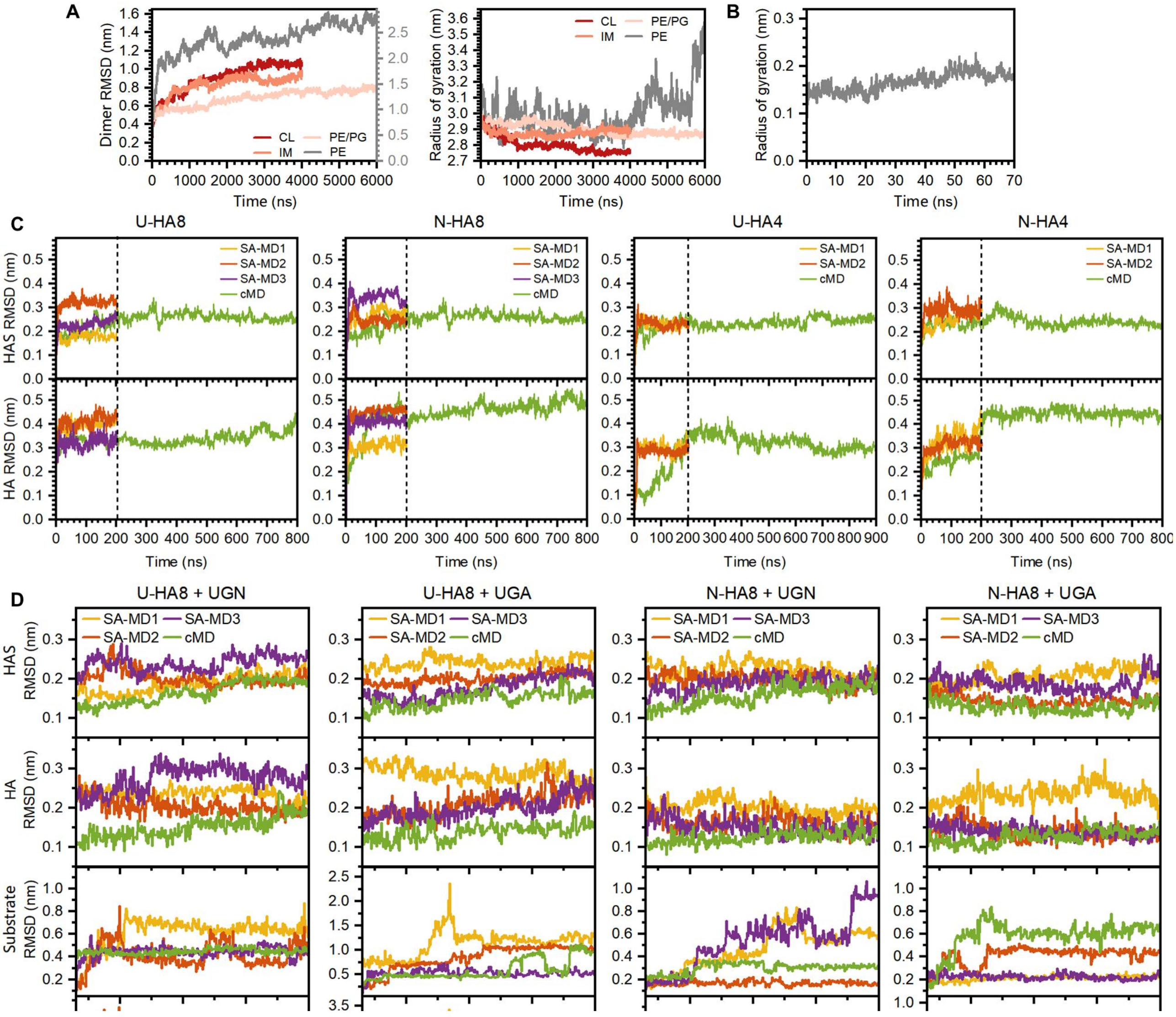


Supplementary Fig. 5 Time profiles of MD simulations. (A) RMSD and radius of gyration for SeHAS dimers during coarse-grained MD. (B) RMSD for SeHAS dimers during all-atom MD. (C) Time profiles for HAS-HA complexes. (D) Time profiles for HAS-HA-substrate complexes, HA8 as examples.
