## Supplementary figures and images for "Substrate tunnel regulates substrate specificity switching in hyaluronan synthase"

### Substrate capture by C-loop.gif

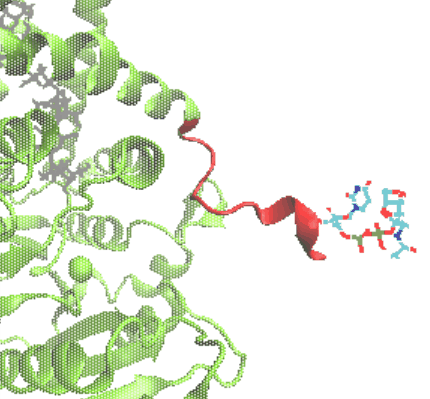

### Substrate retention by C-loop.gif

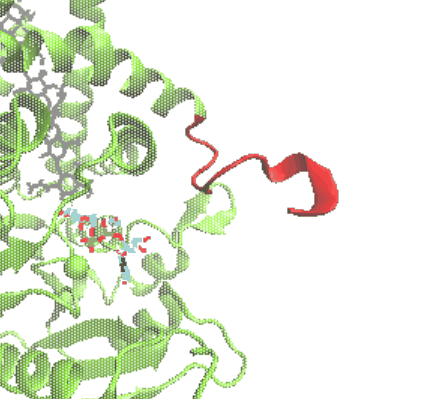
